# Within-Trial Noise Accounts for Inhibition of Return

**DOI:** 10.64898/2026.05.05.722974

**Authors:** Tal Seidel Malkinson, Alexia Bourgeois, Nicolas Wattiez, Ana B. Chica, Pierre Pouget, Paolo Bartolomeo

**Author notes:** These authors contributed equally to this work (Pierre Pouget, Paolo Bartolomeo).

## Abstract

Inhibition of return (IOR) refers to the slowing of response times (RTs) for stimuli presented at previously inspected locations relative to novel locations. However, the exact processing stage(s) at which IOR occurs, and its nature across different response modalities, remain debated. By reanalyzing RT data from a target-target IOR paradigm with a single noisy accumulator model, we tested whether IOR could occur at sensory or attentional stages of processing, or at later stages of decision and action selection. We considered IOR under two conditions: manual and saccadic responses. The within-trial Gaussian noise parameter best explained both manual and saccadic IOR, suggesting that in both modalities, IOR may result from a more fluctuating accumulation of evidence for repeated locations. These results support the hypothesis that target-target IOR may primarily involve attentional-level mechanisms.

**Significance statement:** We respond more slowly to a stimulus that is presented within a short interval in the same location (“inhibition of return”), a bias thought to promote efficient visual exploration. Using evidence-accumulation modeling of manual and eye-movement reaction times from two previous studies, we found that the key change linked to inhibition of return is greater within-trial variability (noise) in evidence accumulation, not a higher decision threshold. Understanding which processing stage is affected can help connect behavioral effects to the brain networks that support attention and orienting.

## 1. Introduction

Inhibition of return (IOR) refers to the slowing of response times (RTs) for stimuli presented at previously inspected locations relative to novel locations (Berlucchi et al., 1981; Klein & Redden, 2018; Lupiáñez et al., 2006; Posner & Cohen, 1984). IOR is believed to promote environmental exploration by avoiding repeated scanning of previously visited locations (Klein, 1988; Klein et al., 2023). This phenomenon has been extensively explored since its discovery in the 1980s (Posner et al., 1985), but its exact nature and neural bases remain debated. Repeated peripheral events can result in faster RTs (RT facilitation) or slower RTs (IOR), depending on several variables, including the delay between the stimuli. This delay varies as function of the motor effector used (manual responses or eye saccades), and the type of visual task (detection or discrimination) (Chica et al., 2014; Lupiáñez, 2010). As noted by some theorists (Berlucchi, 2006; Chica et al., 2014; Lupiáñez, 2010), this pattern of context-dependent outcomes has been argued to sit uneasily with a simple “inhibited attention return” account of IOR, because an invariant attentional inhibition would not straightforwardly predict both facilitation and inhibition across these conditions (Chica et al., 2014; Klein & Redden, 2018; Posner et al., 1985). Moreover, the neural bases of IOR, as well as the specific processing stage(s) involved in IOR generation, remain contentious. IOR could occur at early sensory or attentional stages of processing, at later stages of decision and action control, or in all stages. Originally, Posner et al. (1985) suggested that IOR results from orienting attention to a location and subsequently removing attention from that location, thereby inhibiting reorientation to the originally attended location. However, it has been shown that this attentional account cannot fully explain IOR and that oculomotor programming makes an important contribution to IOR generation (See Klein, 2000 for a review). Converging evidence suggests that IOR could delay both motor responses and the return of exogenous attention (Chica et al., 2010; Lupiáñez et al., 2006).

It has been consistently demonstrated that IOR is not a unitary phenomenon but instead comprises two distinct forms: an input form and an output form. These variants have been extensively described, establishing that the input form affects sensory and perceptual processing when the reflexive oculomotor system is suppressed, whereas the output form reflects a bias in response selection or execution when that system remains active (Chica et al 2010; Taylor & Klein, 2000, Hilchey, Klein & Satel, 2014, Klein & Redden, 2018; Redden et al., 2021). To differentiate between these two types, three fundamental diagnostic tests have been developed. The first is the accuracy diagnostic in speed-accuracy tradeoff-space, which reveals that the output form is characterized by slower but more accurate responses (the hallmark of a speed-accuracy tradeoff), while the input form manifests as slower responding accompanied by a reduced accuracy (Chica et al. 2010). The second diagnostic involves the dissociation between peripheral onsets and central arrow targets, where the output form generates inhibition for both stimulus types due to its directional or motor nature, in contrast to the input form which only inhibits responses to stimuli presented at the physical location previously attended (Taylor & Klein, 2000, Hilchey, Klein & Satel, 2014, Klein & Redden, 2018; Redden et al., 2021). Finally, the locus-of-slack logic within the psychological refractory period (PRP) paradigm demonstrates that the input form operates at early (Kavyani et al., 2017), pre-bottleneck stages of processing, allowing its inhibitory effect to be absorbed into cognitive slack at short intervals, while the output form operates at late, post-bottleneck stages and produces an additive delay regardless of the interval between targets (Klein et al., n.d., 2020).

Several subcortical and cortical regions have been implicated in the generation of IOR. For example, neuropsychological evidence (Sapir et al., 1999) and non-human primate electrophysiology (Dorris et al., 2002) suggest an important role for the midbrain superior colliculus (SC). However, Dorris et al. (2002) demonstrated that the SC is not the productive site of signal reduction. They suggested that signal reduction could potentially be generated in the posterior parietal cortex (PPC). Supporting evidence for this hypothesis comes from electrophysiological studies showing that neural activity in the monkey lateral intraparietal cortex (LIP) is reduced for already explored targets during visual search (Mirpour et al., 2009). Further evidence comes from the observation that in human patients with right hemisphere damage and visual neglect, manual IOR (the input form, requiring the suppression of reflexive eye movements) was reduced for right-sided repeated stimuli (Bartolomeo et al., 1999, 2001), where it could even revert to a “facilitation of return”, i.e., faster RTs for repeated targets (Bourgeois et al., 2012). An advanced lesion analysis study showed facilitation of return in patients with damage to the supramarginal gyrus in the right parietal lobe or to its connections with the ipsilateral prefrontal cortex (Bourgeois et al., 2012). Importantly, however, these patients showed normal saccadic IOR (the output form, requiring reflexive eye movements). Indeed, a contribution of the frontal eye field (FEF) to IOR has also been reported (Mirpour et al. 2019; Ro et al. 2003), and FEF connections with the supramarginal gyrus have been shown to support visual attention shifts (Heinen et al., 2017).

Recently, an intracerebral recording study in drug-refractory epileptic patients (Seidel Malkinson et al., 2024) used a peripheral cueing detection paradigm to establish the critical involvement in the generation of manual IOR of a specific cluster of intracerebral contacts, located at the center of a large-scale cortical gradient extending from the visual cortex to frontoparietal regions. The gradient mapped onto the brain’s core–periphery topography and revealed a shift in neural activity along it, transitioning from visual modulation at the periphery to response-related modulation at the core. Attentional effects emerged at the gradient center, at the intersection of visual and response signals (Seidel Malkinson et al., 2024). Specifically, neural activity in this cluster peaked 22 ms later when the cue and target appeared at the same location than when they appeared at different locations, a pattern mirroring behavioral IOR.

Additional evidence regarding the causal role of cortical structures in IOR generation comes from transcranial magnetic stimulation (TMS) studies (Satel et al., 2019). For instance, Bourgeois et al. (2013a, 2013b) examined how the frontoparietal attention network contributes to manual and saccadic IOR. They found that disrupting the intraparietal sulcus (IPS) and temporo-parietal junction (TPJ) in either hemisphere affected IOR in different ways, depending on the hemifield and the response type. Specifically, TMS over the right TPJ interfered with manual IOR for right-sided targets, while TMS over the right IPS disrupted manual IOR for both hemifields, and saccadic IOR for left-sided targets. Stimulation of the left hemisphere had no effect. These findings were explained by a theoretical model proposed by Seidel Malkinson and Bartolomeo (2018), labeled FORTIOR (Frontoparietal Organization for Response Times in IOR), stipulating the cortical basis of IOR generation in target-target detection paradigms, and accounting for the complex pattern of interference produced by TMS stimulation. Based on the known architecture of frontoparietal cortical networks and on their anatomical and functional asymmetries (Bartolomeo & Seidel Malkinson, 2019), FORTIOR postulates that both manual and saccadic IOR arise from a noise-enhancing reverberation within spatial representations guiding behaviour, implemented in frontoparietal circuits linking frontal eye field (FEF) and intraparietal sulcus (IPS), but likely not restricted to them. The noisier saliency signal is then forwarded to the manual and saccadic motor systems, thereby eliciting IOR. Differences between the readout capacities of the manual and saccadic effector systems when reading the output of the FEF-IPS circuit may then lead to the observed dissociations between saccadic and manual IOR. This account does not imply that IOR depends exclusively on cortical mechanisms, as similar computational principles may also be implemented in subcortical structures such as the superior colliculus.

Klein and Redden (2018) proposed a different view, positing two types of IOR: an ‘input’ form that causes IOR by biasing perception, and an ‘output’ form that causes IOR by biasing action (see also Chica et al., 2010). Specifically, Klein and Redden suggested that input-IOR is generated when the reflexive oculomotor system is actively suppressed (i.e., when participants must suppress reflexive eye movements by maintaining fixation (Chica et al., 2010) or making antisaccades (Redden et al., 2016), and results from modulation of activity in early sensory pathways in retinotopic coordinates, rather than from response outputs. Output-IOR occurs when the reflexive oculomotor system is not suppressed, and involves projections from the SC to cortical areas (i.e., FEF and posterior parietal cortex). Output IOR is spatiotopic and independent of the response modality. Therefore, according to this account, manual IOR (generated while suppressing reflexive eye movements) and saccadic IOR reflect distinct signal-reduction processes and involve distinct underlying neural mechanisms. Klein & Redden’s (2018) distinction is not defined by response modality alone, but by the state of the reflexive oculomotor system and by task features that enforce or relax oculomotor suppression. In particular, Klein Lab studies often implement trial-by-trial feedback to ensure fixation compliance (and thus sustained oculomotor suppression) and use additional diagnostic manipulations (e.g., accuracy/SAT patterns, central vs peripheral targets, PRP) to distinguish perceptual (“input”) from motoric (“output”) contributions. The present study re-analyses a target-target paradigm that includes fixation monitoring/exclusion but does not implement the full set of diagnostic constraints; we use the input/output terminology descriptively and therefore do not claim any definitive classification for manual vs saccadic conditions within the full Klein Lab framework.

Accumulator models, which explain the decision to respond as an accumulation of evidence to a response threshold (see reviews by Luce, 1986; Smith & Ratcliff, 2004), can provide an important contribution to our understanding of IOR. Both behavioural and neurophysiological evidence suggest that the accumulation rate is affected by the stimulus quality and that the evidence criterion is modulated by the prior probability of a particular response (Gold & Shadlen, 2007; Pouget et al., 2011). In the case of IOR, increased RTs for repeated targets might be explained by a variety of model parameters (Satel et al., 2019), such as sensory or attention-level effects (e.g., slower rate of accumulation or decreased noise), or a higher decision-level effect (larger distance between baseline and decision threshold, jointly referred to here as threshold). Identifying the relevant parameters may thus help determine the processes implicated in IOR. Presently, only three studies used this approach to explore IOR. The first modelled RT data from two experiments in which participants generated sequences of three saccades in response to a peripheral or central cue, yielding saccadic IOR for movements to the immediately preceding fixated location (Ludwig et al., 2009). Using a linear ballistic accumulator model, Ludwig et al. (2009) asked whether changes in accumulation rate or in decision threshold best explain saccadic IOR effects. They found that saccadic IOR was best accounted for by a change in the accumulation rate. Ludwig et al. (2009) concluded that an attenuated sensory response to stimulation at a recently fixated location might account for a lower rate of response accumulation at that location. This finding was the first to propose a plausible explanation for a reduced accumulation rate under peripheral cueing conditions. However, the authors did not directly test the potential effect of changes in accumulation noise during the task, nor did they extend their findings to manual responses. Therefore, this study could not provide evidence for such an account in data produced with manual response, which are non-spatially directed (see also Rafal et al., 1994; Taylor & Klein, 2000).

In the second study, MacInnes (2017) modelled data from two experiments testing the gradient of IOR as a function of random, continuous cue-target Euclidean distance and cue-target onset asynchrony. Although behavioral RTs differed across response modalities, best-fit models indicated similar IOR gradient parameters for the two effector types, suggesting similar underlying mechanisms for saccadic and manual IOR. MacInnes (2017) found that the model parameter best explaining location-associated changes in RT was the variance of the accumulation starting point for both manual and saccadic responses. As the spatial distance between the cue and the target increases, the variability in the starting point of information accumulation also increases. This, on average, shifts the accumulation closer to the decision threshold and thus decreases RTs. This result is consistent with neural findings, in which changes in resting-state activation levels in SC (Dorris & Munoz, 1998) depend on target-location probabilities. The divergent results for IOR-associated model parameters in the studies by Ludwig et al. (2009) and MacInnes (2017) may stem from differences in experimental paradigms and model assumptions. In particular, Ludwig et al. examined sequential saccades in a discrete target paradigm using a linear ballistic accumulator model focused on rate and threshold parameters, whereas MacInnes modelled spatial gradients of IOR across continuous cue–target distances using a diffusion framework in which variability in starting point played a central role. These differences make direct comparison of parameter interpretations difficult across studies.

Finally, Redden and colleagues (2020) used a drift-diffusion model to test which parameter best accounts for saccadic IOR in a non-spatial discrimination task with uninformative peripheral cues under two conditions: prosaccades and antisaccades. They found that, in the prosaccade condition, in which the reflexive oculomotor system is not suppressed, IOR was accounted for equally well by the threshold and trial noise parameters (Redden et al., 2020). In the antisaccade condition, in which the reflexive oculomotor system was suppressed, the best-explaining model parameter was the rate parameter (Redden et al., 2020). The rate parameter reflects the speed of evidence accumulation and is commonly interpreted as indexing the quality or strength of the sensory signal driving the decision process. Thus, they concluded that, depending on the state of the oculomotor system, IOR may arise from two distinct processes, which are at odds with the aforementioned MacInnes (2017) model.

Importantly, the previous modeling approaches to inhibition of return (IOR) have differed substantially in their assumptions, parameterizations, and the specific aspects of behaviour they were designed to capture. For example, Ludwig et al. (2009) used a linear ballistic accumulator model, a relatively minimal framework well suited to testing whether IOR effects arise from changes in accumulation rate or decision threshold, but which did not explicitly consider variability in the accumulation process. In contrast, MacInnes (2017) employed diffusion-based models incorporating spatial gradient parameters, allowing the investigation of how IOR varies as a function of spatial distance between successive stimuli. More recently, Redden et al. (2020) used a drift-diffusion framework capable of jointly modeling response times and accuracy, thereby enabling the study of speed–accuracy tradeoffs and providing a richer characterization of decision processes.

Rather than viewing these approaches as contradictory, they can be understood as emphasizing different components of a multidimensional process shaped by differences in task design, dependent measures, and model architecture. In this context, the present study adopts a simplified stochastic accumulator framework to test whether differences between Return and Non-return trials are best accounted for by changes in accumulation rate, decision threshold, or within-trial variability. In contrast to previous work, our approach focuses specifically on within-trial variability and applies a unified modeling framework to both manual and saccadic responses within the same participants, allowing us to test whether a common computational mechanism can account for IOR across response modalities.

To implement this approach, we used a simplified stochastic accumulator (race-type) model derived from evidence-accumulation frameworks (Boucher et al., 2007; see also Noorani, 2014; Schall, 2019, for reviews). Conceptually minimalist and mathematically tractable, this family of models offers a simple account of RTs. A simple, formal view of our version of the stochastic accumulator model is illustrated in Figure 3 and described in Equation 1. An additional constant non-decisional latency, including afferent and efferent delays associated with stimulus encoding and peripheral motor delays, is also added to ‘the decision period’ (Brown & Heathcote, 2008; Pouget et al., 2009). In our model, a decision signal starts from a unique starting point and rises toward a threshold. Once the signal reaches the threshold, a decision is made regarding the appropriate action.

In contrast, the present approach focuses on within-trial variability in the accumulation process and applies a unified modeling framework to both manual and saccadic responses within the same participants. This allows us to test whether a common computational mechanism can account for IOR across response modalities.

Specifically, we had two aims: 1. Determine if IOR RT-distributions could be best accounted for by sensory or attentional parameters (rate of accumulation and noise), or by decisional processes (decision threshold); 2. Test whether the same model parameters can account for RT-distributions for manual and saccadic IOR, and thus reflect a common mechanism (note that our goal was not to directly compare the value of each parameter between the saccadic and motor conditions, but rather to reveal the best explaining parameter within each modality). Finally, since the activity of movement neurons in FEF and SC has been described as instantiating a stochastic accumulation process (Boucher et al., 2007; R. H. S. Carpenter et al., 2009; R. H. Carpenter & Williams, 1995; Hanes & Schall, 1996; Ratcliff et al., 2007; Schall, 2004), we hope that future studies will permit us to investigate the links between model and neurophysiological response.

## 2. Materials and Methods

### 2.1 Experimental data

The RT data used for modeling were obtained from the pre-TMS condition (i.e., excluding TMS-related perturbations) of the studies by Bourgeois et al. (Bourgeois et al., 2013a, 2013b). In these studies, participants performed a target-target paradigm (See Maylor & Hockey, 1985). Four black peripheral circles surrounding a circle placed at fixation were displayed (see Fig. 1). Participants had to respond as fast as possible to one of the peripheral circles becoming white, either by pressing a key while maintaining fixation, or, in a different session, by making a saccade towards the target. In the manual condition, ocular fixation was monitored online (eye tracking) and trials with fixation breaks were excluded, but participants did not receive trial-by-trial feedback contingent on fixation performance. Thus, the manual task strongly encouraged fixation but did not implement the more stringent feedback-based enforcement often used to operationalize reflexive oculomotor suppression in KleinLab paradigms. participants’ compliance with fixation instructions was remarkably high. We note, however, that participants broke fixation in the manual condition in only 1.05% and 3.66% of trials, respectively, in the original experiments by Bourgeois et al. (2013a, 2013b). The behavioral experiment was sponsored by the Inserm Clinical Research Committee (C08-45) and approved by the Île-de-France 1 Institutional Review Board (study no. 2009-A-00035-52). Here, we compared RTs for targets appearing at the same location as the preceding one (Return trials) with RTs for targets appearing at new locations on the diagonally opposite sides of the display (i.e., a bottom-left target after a top-right target; Non-return trials). Responses to right- and left-sided targets were pooled within each return category to maximize modeling robustness. We modelled RT data from 34 participants from the original study (Right TPJ n=8; Right IPS n=8; Left IPS n=8; Left TPJ n=10). The separate control group reported in Bourgeois et al. (2013b) was not included.

**Figure 1.**
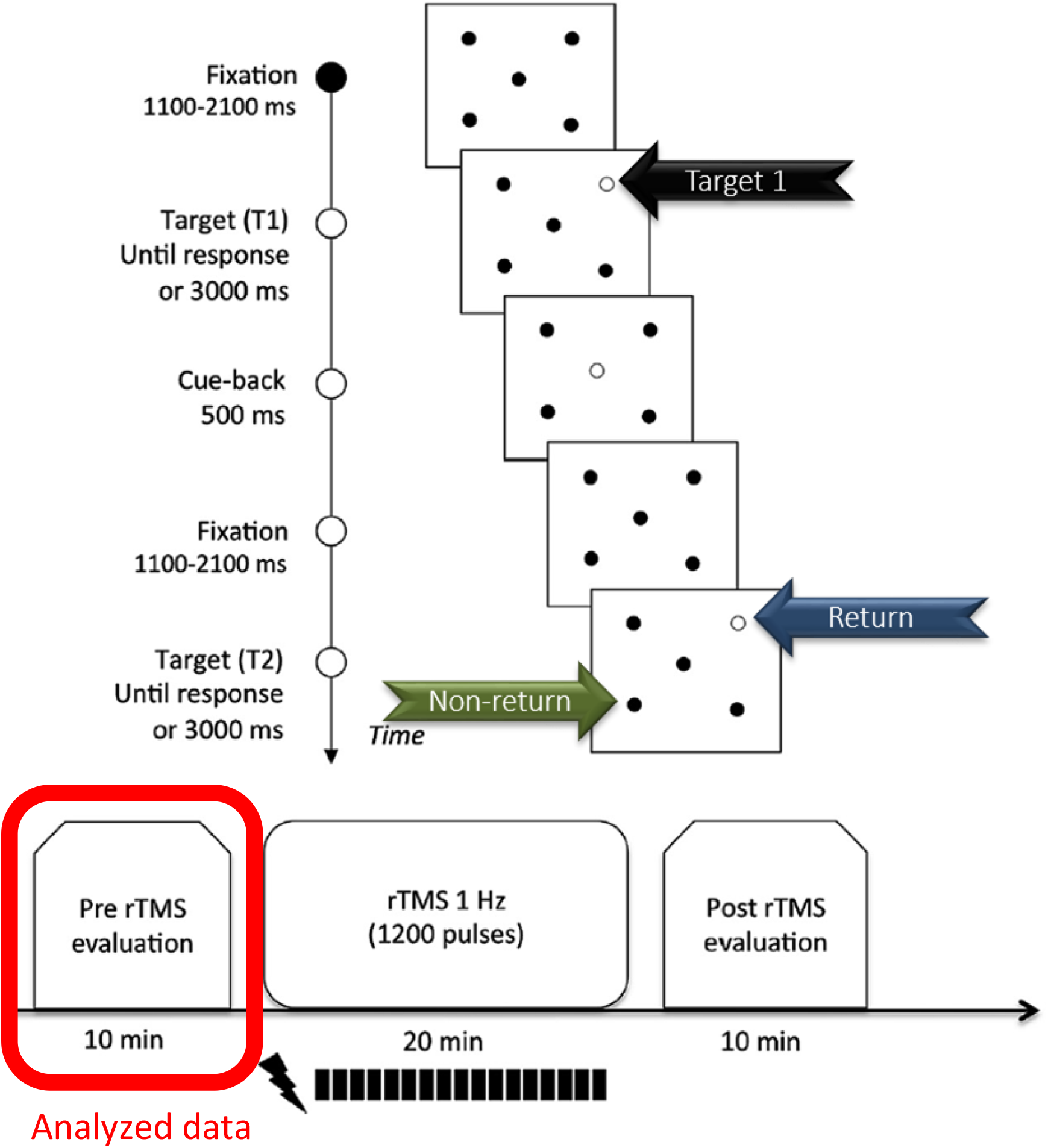
Target-Target detection task in Bourgeois et al., 2013a, 2013b. A. Trial sequence and timing. In the manual task, participants maintained fixation and manually detected the appearance of peripheral targets. In the saccadic task, participants moved their gaze to peripheral targets and back to the center upon cue-back appearance. B. Timeline of the behavioral and Repetitive TMS (rTMS) conditions of the original study. Two runs of each task (manual and saccadic) were performed for each participant in two different session. A 10 min run was performed immediately before (pre-TMS) and the other one immediately after (postTMS) repetitive TMS stimulation. The data analyzed in this study consisted of only the Targets repeating in the same exact location (Return trials, blue arrow) and targets repeating in diagonally opposite side (Non-return trials; green arrow) from the previous target (Target 1; black arrow), only during the pre-TMS run (circled in red). Adapted from

## 2.2 Model fitting

Formally, the go unit accumulates activation according to the following stochastic differential equation (Boucher et al., 2007):

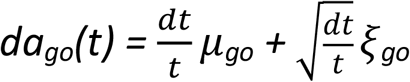

This equation (see Appendix A Supplementary Table A1 for parameter details) specifies the change in unit activation da_go_ within a time step dt (dt/t set equal to 1). The mean growth rates of the go unit are given by the *μ*go. *ξ* is a Gaussian within-trial noise term with a mean of zero and a variance of 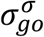. The race ended when the unit crossed the threshold within 3,000 ms. If unit activity is negative during the race, it is reset to zero at this point (i.e., a non-physiological value). An additional 100 ms delay was introduced to account for stimulus encoding within the go unit.

The observed latency distributions for correct responses, as generated by the model, were binned into 5 quantiles, defined by the following boundaries: {0, 0.2, 0.4, 0.6, 0.8, 1.0}. A local chi-square was computed at each bin, and a general chi-square was obtained by summing each local chi-square. To find the best parameters for each latency distribution, we minimized the general chi-square by using a minimization function (patternsearch from the global optimization toolbox of Matlab). Patternsearch searches for a minimum using an adaptive mesh aligned with the coordinate directions. Because minimization functions are sensitive to their starting points, we conducted pattern search from 50 randomly selected starting points. Finally, to avoid a local minimum, we again started patternsearch from 200 new start points. These starting points were determined by the best parameters from the first run and were defined as follows: best parameters ± 0.01 units and best parameters ± 0.02 units. After applying this procedure to each distribution of the Return and the Non-Return conditions, we ran a new procedure in order to determine which of the *μ, σ*, or *θ* parameters could explain the differences observed in the Return and the Non-Return latency distributions. To do this, we fixed two parameters of the Return condition on the latency distribution of the Non-return condition and let one parameter free. As in the first procedure, pattern search was run from 50 randomly selected starting points, followed by 200 additional starting points. All these procedures were performed on a supercomputer cluster (NEC, 40 nodes, 28 CPU Intel Xeon E5-2680 V4 2.4 GHz/node, 128 GB RAM/node).

To facilitate interpretation, we relate the model parameters to broad classes of cognitive processes. The accumulation rate is typically associated with the quality of sensory evidence and perceptual processing, whereas the decision threshold reflects the amount of evidence required before committing to a response and is therefore linked to decisional or strategic factors. The within-trial variability parameter captures fluctuations in the accumulation process and may reflect variability in attentional or priority-related signals that influence the stability of evidence accumulation. These mappings are not process-pure, and each parameter may be influenced by multiple underlying mechanisms. Accordingly, parameter interpretations should be understood as reflecting dominant contributions rather than exclusive mappings.

The analysis code can be obtained by request from the corresponding author.

### 2.3 Statistical analysis

Following (Smith & Ratcliff, 2004) (2004) the optimized fit of each model can be minimized using (χ^2^) statistics (Pouget et al. 2011). However, to account for the number of parameters as well as the variance, here we used Bayesian information criterion (BIC) values that were computed for each model using Matlab (MATLAB, 2017) and served for model estimation and comparison. While BIC allows identification of the best-fitting parameter among competing models, it does not imply that the remaining parameters do not contribute to explaining the observed data. RTs and BIC values were compared statistically using JASP ((Jasp Team, 2020, JASP (Version 0.12)[Computer software]). We modelled RT data from 34 participants. To control for inter- and intra-subject RT variability, and in accordance with the procedure in Bourgeois et al. (2013a, 2013b), trials with RTs faster than 100ms (anticipations) and trials with RTs slower than 2.5 SD were excluded. For RT analysis, the same trial-selection criteria were applied across the 34 participants. However, the present trial selection procedure differs from that originally used by Bourgeois et al. (2013a, 2013b) in that it pooled data for right- and left-sided targets to maximize the number of modelled trials. When possible, repeated-measures ANOVAs (with Greenhouse-Geisser correction when necessary) and Holm-corrected post hoc tests were used to analyze RT and model parameters. BIC values did not deviate significantly from a normal distribution (Shapiro-Wilk test, Return *µ*-free, *σ*-free and *θ*-free models: W=0.960, *p*=0.81; W=0.930, *p*=0.51; W=0.962, *p*=0.83; respectively; Non-return *µ*-free, *σ*-free and *θ*-free models: W=0.948, *p*=0.67; W=0.933, *p*=0.54; W=0.975, *p*=0.93, respectively). Therefore, repeated measures ANOVA was used to test the main effects of Return/Non-return on RTs and BIC values. Descriptive statistics are reported as *X* ± *SE*.

No part of the study procedures or analyses was pre-registered prior to the research.

## 3. Results

### 3.1 Behavior

Across participants, after excluding anticipatory responses and responses slower than 2.5 SD, the median number of included trials for manual responses in the Return and Non-return conditions was 30 (range, 14-49) and 33 for saccadic responses in Return and Non-return conditions (range, 20-72).

Fig. 2 shows the mean saccade latencies and manual response times across subjects, for each of the conditions of Return vs. Non-return (see Appendix A Supplementary Figures A1-2 for individual reciprobit plots in the Manual and Saccadic conditions). A number of results emerge. First, a two-factor repeated-measures ANOVA was performed to assess the reliability of the Return effect across manual and saccadic modalities. The difference between Return versus Non-return trials was significant (Return vs. Non-return: 287±6 ms vs. 273±6 ms; F_(1,30)_=51.34, *p*<0.001, ⍰^2^=0.03; see Figure 2). There were also significant main effects of Modality (Manual vs. Saccadic: 304±11 ms vs. 256±4 ms; F_(1,30)_=19.70, *p*<0.001, **η**^2^=0.37) and a significant interaction between Return and Modality (Manual IOR vs. Saccadic IOR: 9.78±3.1 ms vs. 19.5±2.7 ms; F_(1,30)_=5.63, *p*=0.024, ⍰^2^=0.004), suggesting that IOR was larger in the saccadic modality, although both IOR effects were statistically significant (post hoc tests Return vs. Non-return: Saccadic IOR *p*<0.001; Manual IOR *p*=0.004).

**Figure 2.**
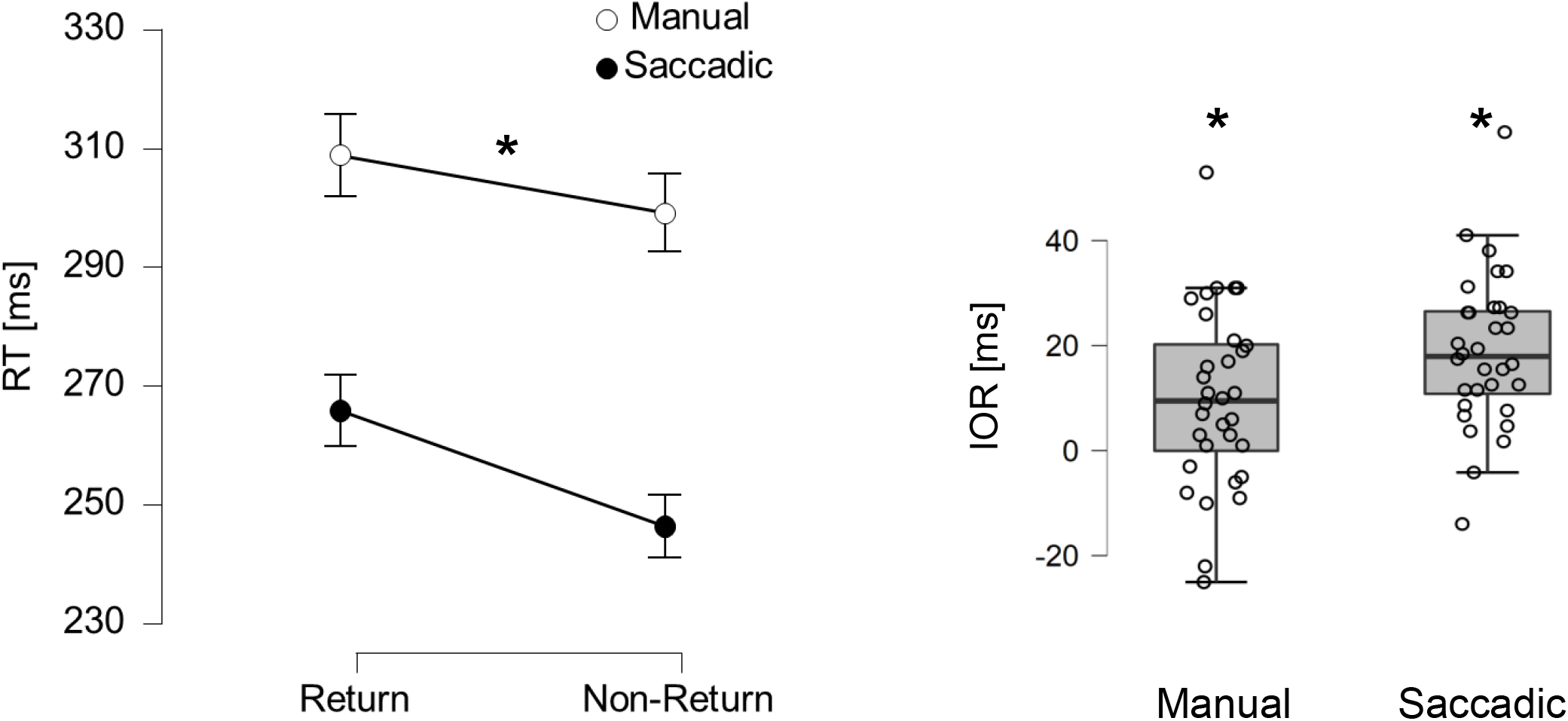
Response time inhibition of return (IOR) effect. Left: Manual response time (RT, white circles) and Saccadic RT (black circles) are longer in Return trials than in Non-return trials (Return main effect, p<0.001). Error bars represent SE. Right: Box plots of the manual (left) and saccadic (right) IOR effects (defined as mean RT_Return_ - mean RT_Non-return_) across participants (circles). Manual and saccadic IOR effects were significant (p=0.004; p<0.001; respectively).

**Figure 3.**
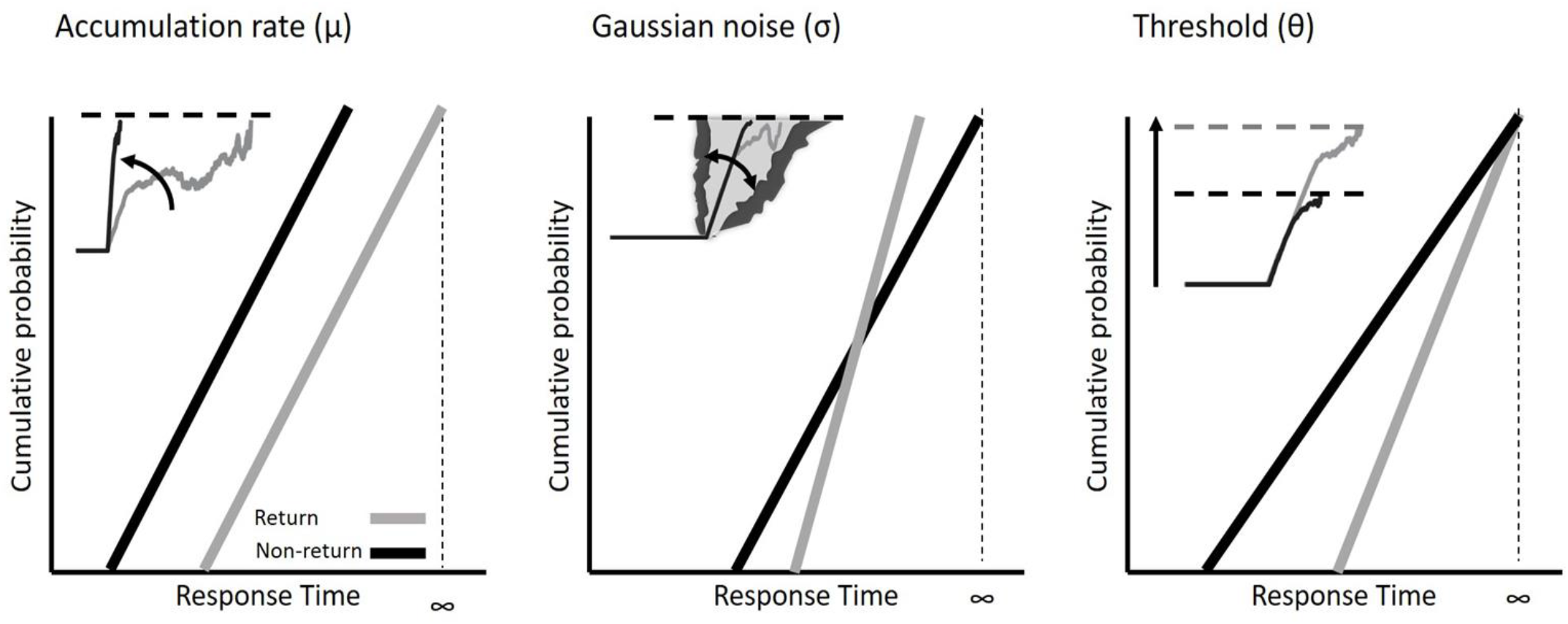
Schematic illustrations of RT distributions for Non-return (black lines) and Return (grey lines) trials. Inset illustrates the simulated model parameter accounting for response time differences due to IOR. The growth rate within the unit, at each time step, is given by a mean in addition to a Gaussian noise term. The estimated RT is given by the time when the unit crossed a threshold within a limit of 3000 ms. In these illustrations, the accumulator coding the Return condition is shown by the grey line. The black accumulators code the Non-return condition. Insets are also schematic illustrations. (Left panel) The target is at a Return location, causing a reduction in the accumulation rate of the associated movement program. As a result, the threshold is crossed later. (Middle panel) The target is at a Return location (grey dots), resulting in a selectively increased within-trial variability in the accumulation rate of the accumulator coding the target and a delay in the time at which the threshold is reached. Note that in this Target-Target task IOR is determined by trial sequence effects and not by within-trial events.(Right panel) The target is at a Return location, resulting in a selective increase in the evidence criterion for the accumulator coding the target and a delay in the time at which the threshold is reached.

### 3.2 modeling

Figure 3 shows illustrative examples of reciprobit plots corresponding to the log-log cumulative probability distribution of saccadic or manual RTs for the Return and Non-return conditions. The figure illustrates examples of single sessions showing RT variations best fitted with a single Manual Saccadic parameter adjustment (*µ, σ*, and *θ*). If IOR is mediated by a change in the underlying rate of accumulation (*µ)*, the rate should be lower for Return responses compared to Non-return responses. If IOR reflects the need for more evidence, the threshold (*θ*) should be raised for Return responses. If changes in within-trial variability of the accumulation rate (*σ*, termed hereafter Gaussian noise) underlie IOR, then the mean accumulation rate for Return responses should be, on average, lower than that for Non-return responses. Note that because we used a target-target task here, the IOR effect is inherently determined by the trial’s immediate history, i.e., the relative location of the target in the previous trial. Therefore, RT changes with successive stimulus presentations, which have previously been used as a measure of the effect of trial sequence (Pouget et al., 2009), are captured here by the *σ* parameter.

#### 3.2.1 The Gaussian noise parameter σ explains manual and saccadic IOR

RTs of Return trials and Non-return trials were separately fitted with accumulator models with 3 free parameters. The resulting Return condition model parameters were used to constrain the Non-return model fit. BIC values were computed for each of the Non-return single-free parameter models, allowing to explore which parameter best accounted for the differences between Return and Non-return trials caused by IOR. This procedure was performed separately for manual and saccadic RTs.

In order to test which parameter: *µ, σ* or *θ*, better explained the differences between Return and Non-return conditions across modalities, we conducted a 2-way repeated measures ANOVA with Free model parameter (three levels: *µ*-free, *σ*-free, or *θ*-free models) and Modality (Manual or Saccadic) as factors. A significant main Free model parameter effect (*µ* BIC: 13.36±0.60, *σ* BIC: 12.17±0.45, *θ* BIC: 13.35±0.54; F_(1.167,37.16)_=8.44, *p*>0.001, η^2^=0.023; see Figure 4), resulting from significant differences between the BIC of the σ-free and the BICs of the *µ*-free and *θ*-free models (Post Hoc comparisons: *µ*-free vs. *σ*-free *p*_*holm*_=0.011; *σ*-free vs. *θ*-free *p*_*holm*_=0.002). There was also a significant main effect of Modality (Manual vs. Saccadic: 14.58±0.55 vs. 11.35±0.82; F_(1,32)_=10.71, *p*=0.003, η^2^=0.188), in which saccadic BIC values were smaller than manual BIC values. The interaction between Modality and Free model parameter factors was not significant (F_(1.404,44.94)_=1.65, *p*=0.2). A follow-up Bayesian model comparison revealed very strong evidence in favor of the full model including the interaction term relative to the null model (BF_10_= 47.98). Importantly, compared with the model including only the main effects of Modality and the Free model parameter (BF_10_= 2.10), the substantial increase in evidence is attributable solely to the inclusion of the interaction term. However, when assessing the interaction across the broader model space using inclusion Bayes factors, the evidence for the interaction effect remained only anecdotal (BF_incl = 2.14). These results suggest that variability in the rate of accumulation best explains RT differences between Return and Non-return trials across both modalities, thereby supporting a perceptual/attentional account of IOR in both manual and saccadic modalities.

**Figure 4.**
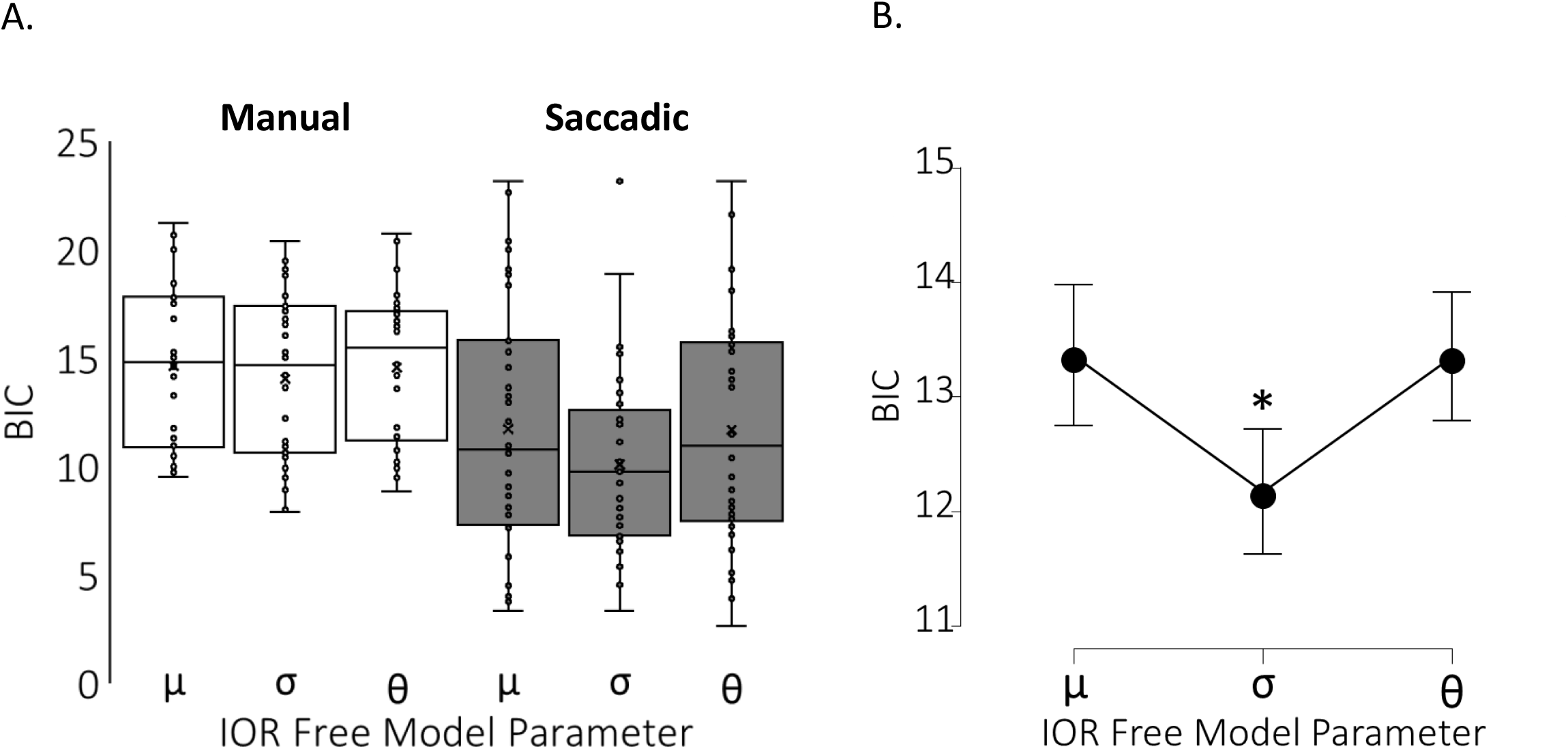
Gaussian noise parameter best explains IOR effect. A. Boxplots of BIC values for µ, σ & θ free models of manual (light) and saccadic (dark) IOR. B. BIC value of the gaussian noise parameter (σ) is significantly the smallest (Free model parameter main effect, p<0.001; Post hoc tests: µ,σ pholm=0.011; θ,σ pholm=0.002). Error bars represent SE.

## 4. Discussion

Does IOR attenuate processes at early sensory or attentional stages, or does it influence later stages such as motor programming? Behavioral evidence suggests that the sensory attenuation of stimulus-related responses may be automatic and non-motor, because it occurs independently of whether an eye or hand response is required, or whether the validity of the cues elicits shorter or longer reaction times (See, e.g., Lupiáñez et al., 2004; Posner & Cohen, 1984). Studies on the neurophysiological substrate of IOR in humans and non-human primates have led to similar conclusions. Reduction of sensory cortical responses has been observed using an event-related potential IOR task (McDonald et al., 1999) and in response to repeated stimuli using a non-RT, delayed match to sample tasks, both in the PPC (Steinmetz et al., 1994; Steinmetz & Constantinidis, 1995) and in the inferior temporal cortex (Miller et al., 1991). This suggests that it is a ubiquitous response property across various brain areas. In the oculomotor subcortical system, and in SC in particular, neuronal recordings have shown correlations between reductions in early sensory neuronal activity in SC and IOR behavior across experimental sessions and on a trial-by-trial basis within a single session (Dorris et al., 2002). All these reduced stimulus-related responses are compatible with a sensory or attentional account of IOR. Despite these results, there remains debate over whether additional motor components contribute to IOR (Ivanoff & Klein, 2001; Taylor & Klein, 2000). Mechanistically, a reduction in the signal-to-noise ratio of the sensory response applied to previously attended objects or regions of a scene is one way to explain IOR. This process would ensure that an initially salient region of the scene might not maintain this status over time. Neurophysiologically, across multiple primary sensory areas, when a sudden stimulus appears at a position outside the receptive field (RF), lateral inhibition dominates the responses inside the RF (Pouget et al., 2005; Schall & Hanes, 1993). When sequential stimulations are presented, RF responses are modulated by lateral inhibition. Specifically, the lateral inhibition following the first stimulation exceeds the termination of the stimulus by about 200ms, and is therefore suitable not only for stimulus competition but also for supporting attentional capture (Lev-Ari et al., 2020). Despite this evidence, our understanding of the complete mechanisms and neurophysiological substrates of IOR remains sparse and fragmentary (Dorris et al., 2002).

Here, we addressed this mechanistic question by employing a modeling strategy that linked RTs to mathematical parameters simulating the various stages of neural processes leading to manual or saccadic responses (Boucher et al., 2007; R. H. Carpenter & Williams, 1995; Gold & Shadlen, 2007; Pouget et al., 2009). This model is a simplified accumulator framework and does not incorporate biologically detailed mechanisms such as leakage or lateral inhibition, allowing us to focus specifically on the contribution of accumulation rate, threshold, and within-trial variability. By examining the rate of evidence accumulation toward a decision threshold, our results indicate that IOR, whether elicited by manual or saccadic responses, primarily depends on the modulation of evidence-accumulation variability. Notably, in our case, the stimulus (a black disk becoming white) had little perceptual ambiguity, which should minimize potential differences in target encoding time between Return and Non-Return trials. Thus, the observed differences should primarily reflect attentional or motor factors. This finding converges with (i) the rTMS effect on right - but not left-posterior parietal cortex reported by Bourgeois et al. (2013a and 2013b) and (ii) IOR-related broadband gamma differences observed in intracerebral recordings (Seidel Malkinson et al., 2024), both pointing to an attentional contribution to IOR. More broadly, it dovetails with MEG evidence that frequency-specific reconfigurations of right-lateralized attentional networks during the cue–target interval predict whether near-threshold targets will be consciously reported (Spagna et al., 2026).

In particular, the intracerebral evidence collected by Seidel Malkinson et al. (2024) in patients with drug-resistant epilepsy, while they were performing a cue-target Posner paradigm, established the critical involvement of a specific cluster of intracerebral contacts (Cluster 2 in (Seidel Malkinson et al., 2024)) in the neural processes underlying IOR. In Cluster 2, neural activity was delayed by 22 ms when the cue and target were presented at the same location, compared to when they appeared at different locations—a timing profile consistent with behavioral IOR. This cluster partially overlaps with the right-hemisphere SLF-II circuit (Bartolomeo et al., 2025), but, as noted by Wang and Theeuwes (2024), it also encompasses contacts in the posterior temporal lobe, including the inferotemporal regions involved in visual recognition.

A key question concerns the interpretation of the within-trial variability parameter (σ). Although this parameter does not map uniquely onto a single cognitive process, its modulation can be understood within frameworks that conceptualize attention as the dynamic allocation of priority across competing representations. In such accounts, attention biases competitive interactions within spatial priority maps, thereby shaping the accumulation of evidence during decision formation (Bisley & Goldberg, 2010; Gold & Shadlen, 2007). Increased variability in accumulation may therefore reflect a less stable or more fluctuating priority signal at previously attended locations. Behaviorally, such increases in within-trial variability are expected to result in broader response time distributions and greater trial-to-trial variability, rather than a simple shift in mean response time. This interpretation is also consistent with the FORTIOR model, which proposes that IOR arises from noise-enhancing reverberation within frontoparietal priority maps. Within this framework, the σ parameter can be viewed as capturing the behavioral consequence of such variability in priority-map dynamics, thereby linking the present computational findings to a broader attentional account of IOR. Although alternative interpretations, including decisional or motor contributions, cannot be fully excluded, this account provides the most parsimonious explanation of the present findings.

The present findings should also be interpreted in the broader context of previous modeling approaches to IOR. Rather than indicating a single definitive mechanism, the diversity of modeling results in the literature likely reflects differences in experimental paradigms, dependent variables, and model assumptions. For instance, Ludwig et al. (2009) reported that saccadic IOR was best explained by changes in accumulation rate, whereas MacInnes (2017) emphasized variability in starting point as a function of spatial distance, and Redden et al. (2020) showed that different parameters (threshold, noise, or rate) may account for IOR depending on task demands and oculomotor state.

In this context, our finding that within-trial variability (σ) provides the best account of RT differences in a target-target paradigm should be understood as complementary to these previous results. Specifically, it suggests that variability in the accumulation process may play a central role under conditions where trial history is a key determinant of behaviour. More generally, these results highlight that different modeling frameworks may capture distinct aspects of IOR, and that apparent discrepancies across studies may arise from differences in task structure and modeling choices rather than fundamental theoretical disagreement.

The ability of a simple stochastic accumulator model to account for IOR effects in a quantitative manner may have important implications. Altered IOR may occur after central nervous system damage (Bartolomeo et al., 1999; Bourgeois et al., 2012; Sapir et al., 1999; Vivas et al., 2006), in psychiatric conditions (Mushquash et al., 2012), or upon neuromodulation of specific brain areas (Bourgeois et al., 2013a; Chica et al., 2011). Examining the underlying parameters of RT distributions may provide novel insights into these deficits and brain functions, rather than simply reporting IOR amplitude or time of occurrence, and can reasonably be expected to reflect the underlying neural processes more directly. In the oculomotor domain, saccade-related neurons in FEF, SC, and Lateral Intraparietal cortex (LIP) are thought to be responsible for sensori-motor transformation (Coles, 1997; Everling & Munoz, 2000; Purcell et al., 2012; Schall, 2019), and may therefore be relevant nodes of the IOR network. A strength of our model is its ability to predict IOR behavior. It represents a simple and robust way of conceptualizing IOR in the specific contexts of target-target paradigms.

In Klein & Redden’s (2018) framework, “input” and “output” IOR are distinguished by the state of the reflexive oculomotor system and by diagnostic task constraints. In our target–target dataset, we observed that the same accumulator parameter (within-trial noise) best accounted for Return vs Non-return in both manual (fixation) and saccadic conditions. Within this paradigm, these results do not support a strong one-to-one mapping between response modality (manual vs saccadic) and distinct underlying mechanisms (input vs output). However, because our dataset does not include the full diagnostic manipulations used to adjudicate input vs output IOR (e.g., feedback-enforced oculomotor suppression; peripheral vs central targets; PRP/SAT), we do not treat the present findings as a definitive test of the broader KleinLab taxonomy. However, while we agree that trial-by-trial feedback is a valuable tool for discouraging eye movements (Hilchey et al., 2014), we remain convinced that the current data set, from the experiments of Bourgeois et al. (2013a, 2013b), provide a robust setting for generating the input form of IOR. Specifically, the Bourgeois et al.’s studies employed a within-participants design where the same individuals performed both manual detection and saccadic localization tasks in separate sessions. This design likely allowed participants to establish and maintain stable and distinct task sets, requiring either the suppression or the activation of reflexive eye movements. Furthermore, participant compliance with fixation instructions was remarkably high, as reflexive eye movements occurred in only <4% of trials in the manual conditions. This extremely low error rate suggests that participants successfully and consistently maintained a mental set of reflexive oculomotor suppression. Consequently, we believe that the resulting IOR was representative of the input form despite the absence of explicit trial-by-trial feedback. Future work that combines feedback-enforced fixation (or antisaccades), central/peripheral targets, and full SAT/PRP diagnostics within the same modeling framework will be needed to directly arbitrate input vs output IOR mechanisms.

Our results appear inconsistent with Klein and Redden’s (2018) modeling results, which identified different explanatory parameters for IOR generated by prosaccades and antisaccades (Redden et al., 2020). Our current evidence, obtained by modeling the results of a target-target paradigm with either manual or saccadic responses, suggests that both manual and saccadic IOR primarily depend on changes in the dynamics of evidence accumulation, with within-trial variability providing the best account in the present data, whose likely neural correlates involve fronto-parietal networks crucial for attentional orienting. In the oculomotor domain, saccade-related neurons in FEF, SC, and LIP are thought to be responsible for the sensori-motor transformation (Coles, 1997; Everling & Munoz, 2000; Mushquash et al., 2012; Schall, 2019), and were shown to be relevant nodes of the IOR network (Mirpour et al., 2009, 2019).

The involvement of these cortical regions in both saccadic and manual IOR in a target-target task was highlighted in the FORTIOR theoretical model (Seidel Malkinson & Bartolomeo, 2018). FORTIOR makes two predictions relevant to the present results. The first postulates that both manual and saccadic IOR arise from reverberation of noise-enhancing activity within priority maps of the frontoparietal circuit linking FEF and IPS. The noisier saliency output is then read by the manual or saccadic motor systems, thereby eliciting IOR. Our finding that the Gaussian noise parameter best explains IOR across both modalities dovetails with this prediction. FORTIOR predicted increased noise in the priority map in Return trials as compared with Non-return trials, whereas the present model captures differences in variability at the level of the accumulation process, which may not map directly onto the form of noise described in FORTIOR. The potential contradiction with our findings could result from different sources of noise in FORTIOR and in our model. This discrepancy highlights, on the one hand, the need to refine and constrain FORTIOR and, on the other hand, the necessity of validating our modeling results with direct electrophysiological evidence.

A second prediction of FORTIOR is that differences between the readout capacities of the manual and saccadic effector systems when reading the output of the FEF-IPS circuit may lead to dissociations between saccadic and manual IOR. For example, the saccade system seems to be more encapsulated (i.e., less prone to interference and illusions) than the manual response system, and relies on a representation that accumulates visual information and location errors over shorter time windows than the representation used for controlling hand movements (Lisi & Cavanagh, 2015, 2017). Our finding that all parameter BIC values were higher in the manual modality than in the saccadic modality is consistent with FORTIOR’s prediction of greater readout reliability for the saccadic system.

It is important to note that our model assumes a constant non-decision time (Da_go_). This assumption might be less accurate for manual responses than for saccadic responses. Another possibility is that different tasks, such as Cue-Target tasks, might reveal the contribution of additional parameters to IOR generation (Bompas et al., 2017; Patel et al., 2010; Pouget et al., 2011). However, remodeling the data with the Da_go_ as a free parameter did not provide significant evidence for Da_go_modality differences. Indeed, task variations are a key difference between the present study and that by Redden et al. (2020), which used antisaccades to inhibit the oculomotor system. However, antisaccade paradigms involve additional control processes related to response inhibition and vector transformation (Munoz & Everling, 2004), which differ from the task demands of the present paradigm. Thus, inhibiting the oculomotor system by enforcing fixation, as in the manual IOR condition used here, is an alternative experimental manipulation with different task demands for testing the effect of the oculomotor system state on IOR. These experimental manipulations may explain the discrepancy between Redden et al.’s (2020) results and those reported here. The question remains open of whether the present model of target-target IOR can be generalized to IOR in cue-target paradigms or in visual search. This is an empirical issue that warrants attention in future research. For the moment, however, we note that there is no principled reason to hypothesize that target-target IOR is a totally different phenomenon from cue-target IOR. For example, both forms of IOR show similar time courses (See Maylor & Hockey, 1985; Posner & Cohen, 1984), and respond in similar ways to brain damage in the right hemisphere (Bartolomeo et al., 1999, 2001; Bourgeois et al., 2012; Vivas et al., 2006). If anything, one might speculate that cue-target IOR, which does not require a motor response to the cue, might be even less sensitive to motor factors than target-target IOR, which requires two successive motor responses.

These task differences also lead to different choices of the accumulation model used. IOR is determined by the location of the target in the current trial relative to that in the previous trial, and not by any sequence of events within the current trial. Nevertheless, the within-trial noise parameter *σ* used here captures the effects of trial sequence (Pouget et al., 2009). Thus, although formally different from the inter-trial noise parameter used by MacInnes (2017), the within-trial noise parameter employed here captures a similar source of variability.

Variability between individuals is a common observation in IOR effect in humans and animals, and also occurred in the group of human participants tested here. The robustness of modeling approaches also depends on the amount of data modelled (the number of participants and input trials). This had limited the subtlety of the effects we could study (e.g., by collapsing left- and right-sided target conditions).

The present findings should be interpreted in light of the dataset used. A strength of the Bourgeois et al. data is the within-participant comparison of manual and saccadic responses in a controlled target–target paradigm. However, the dataset was not designed to dissociate different forms of IOR, as it lacks key diagnostic manipulations such as trial-by-trial feedback on eye movements and detailed accuracy measures. Future studies combining modeling with paradigms that explicitly manipulate oculomotor state and include accuracy constraints would provide a more direct test of the model’s predictions.

The enthusiasm for the neurophysiological approach to tracking the IOR effect is somewhat difficult to reconcile with the small changes in response times observed in IOR manipulations (But see Dorris et al., 2002). The relatively small magnitude of the IOR effect and its inter-individual variability limit the robustness of modeling approaches that aim to elucidate it. To make it truly useful, the approach would ideally benefit from modeling hypotheses that are strong enough to be tested, rejected, or validated. Further human intracerebral studies, including single-neuron recordings from parietal and prefrontal regions, are needed to clarify the precise dynamics and network architecture underlying IOR generation.

## Author Contributions

Conceptualization, T.S.M, P.P., and P.B.; methodology, T.S.M, N.W., P.P; software, N.W.; formal analysis, T.S.M. and N.W.; investigation, A.B., A.B.C.; resources, A.B, P.B., A.B.C. and P.P.; data curation, A.B. and A.B.C.; writing—original draft preparation, T.S.M.; writing—review and editing, T.S.M, A.B., N.W., P.P. A.B.C and P.B.; visualization, T.S.M.; supervision, P.P. and P.B.; funding acquisition, T.S.M and P.B. All authors have read and agreed to the published version of the manuscript.

## Institutional Review Board Statement

The behavioral experiment was sponsored by the Inserm Clinical Research Committee (C08-45) and approved by the Île-de-France 1 Institutional Review Board (study no. 2009-A-00035-52).

## Data Availability Statement

Data and analysis code can be obtained by request to the corresponding author.

## Appendix A

**Supplementary Figure A1.**
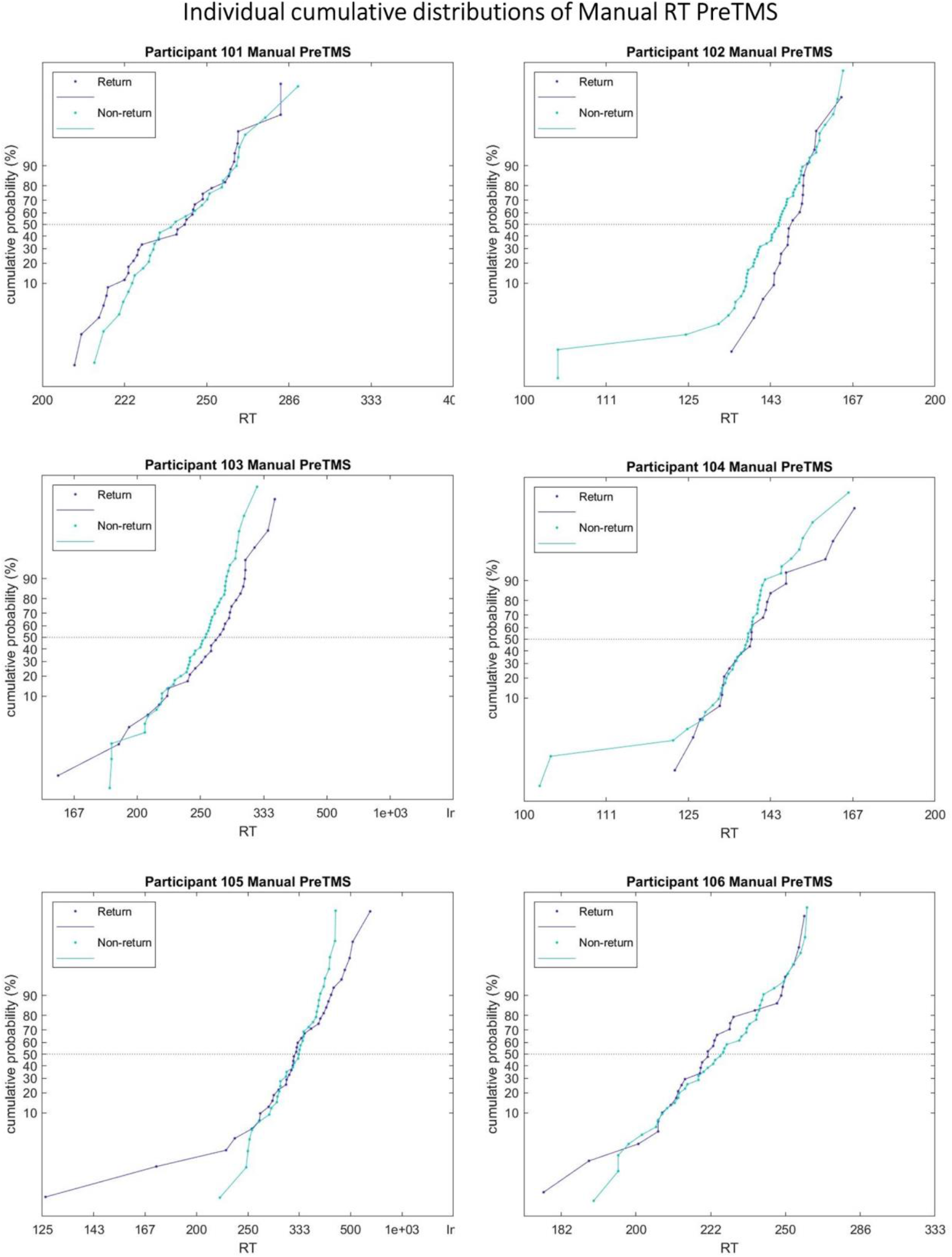
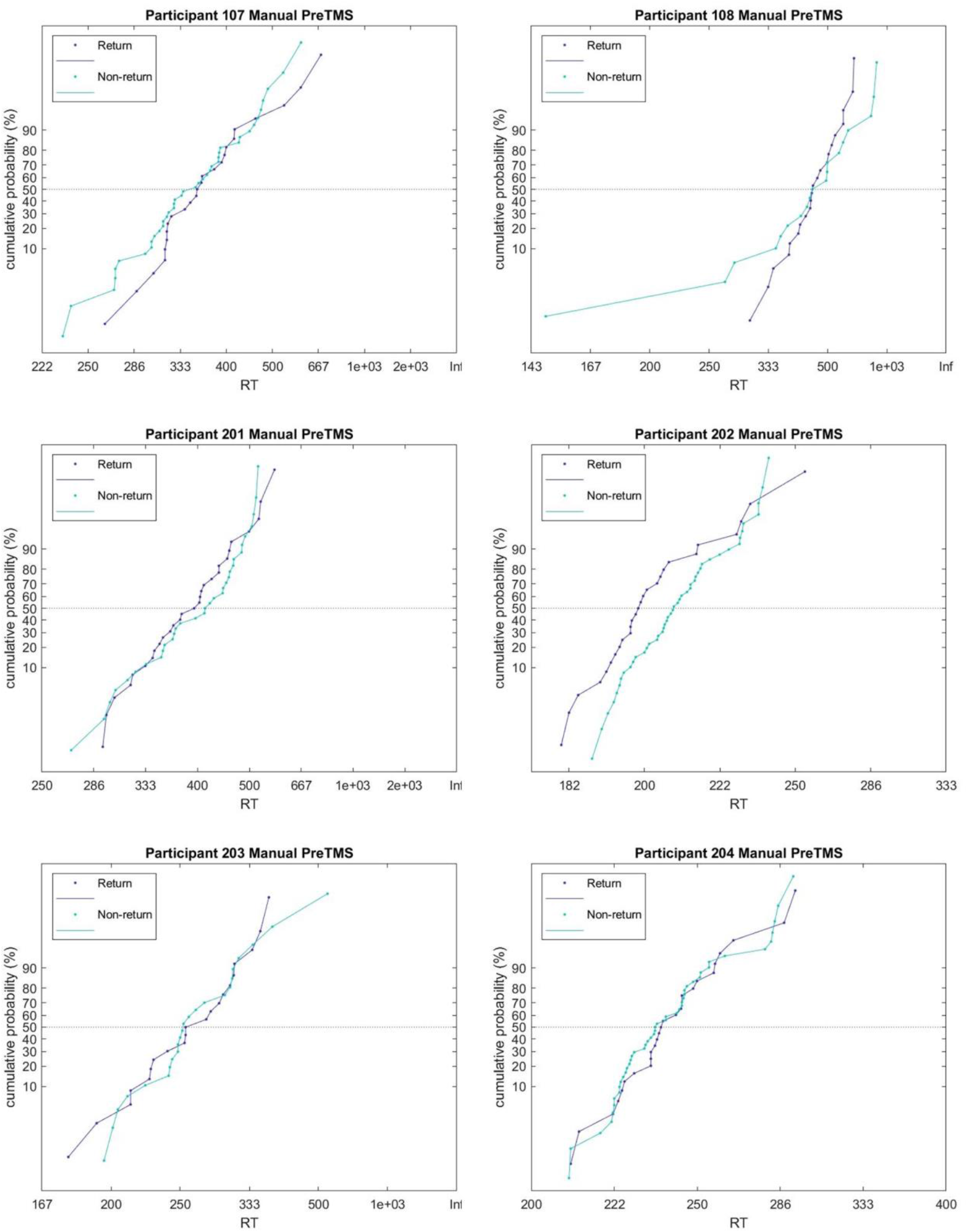
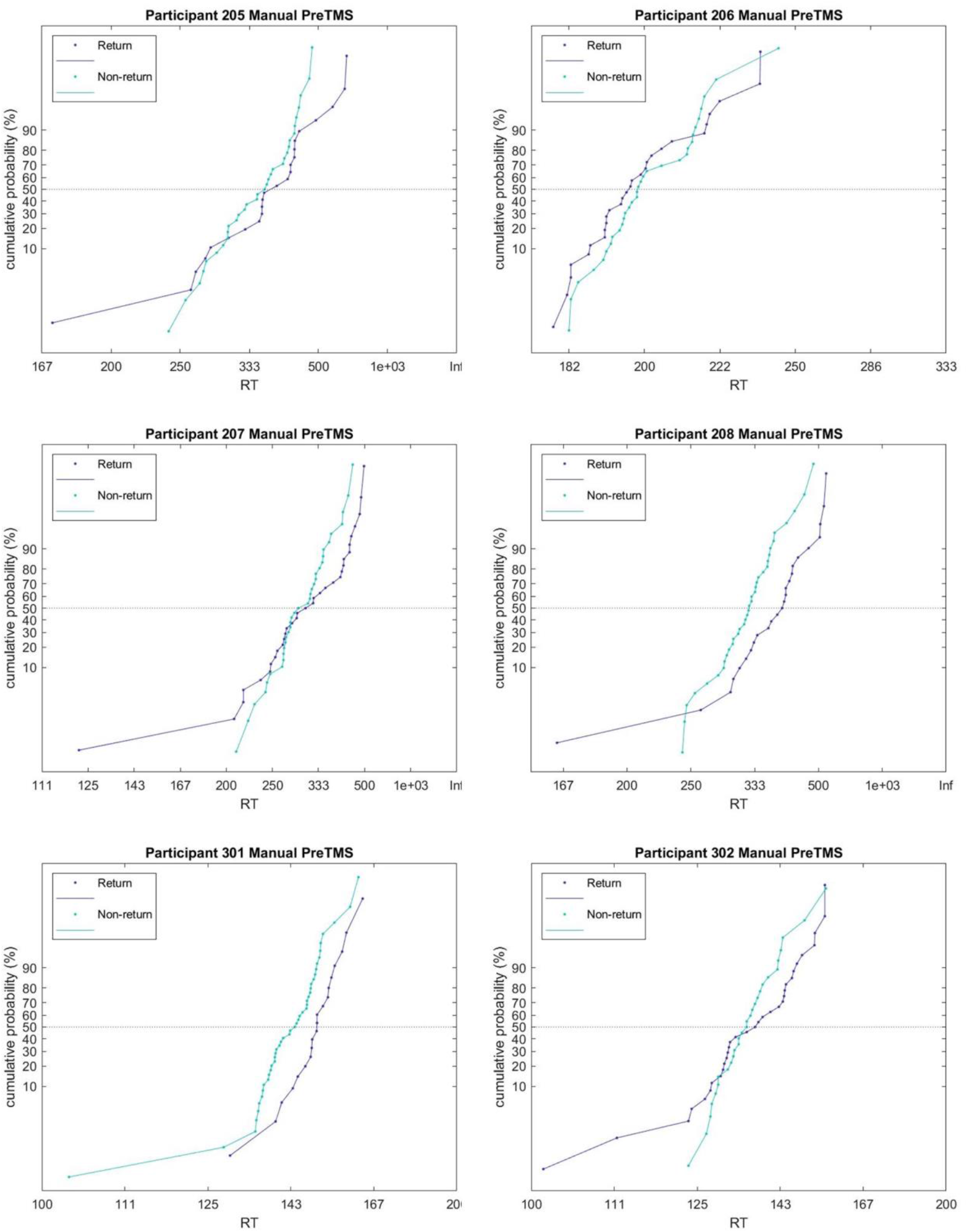
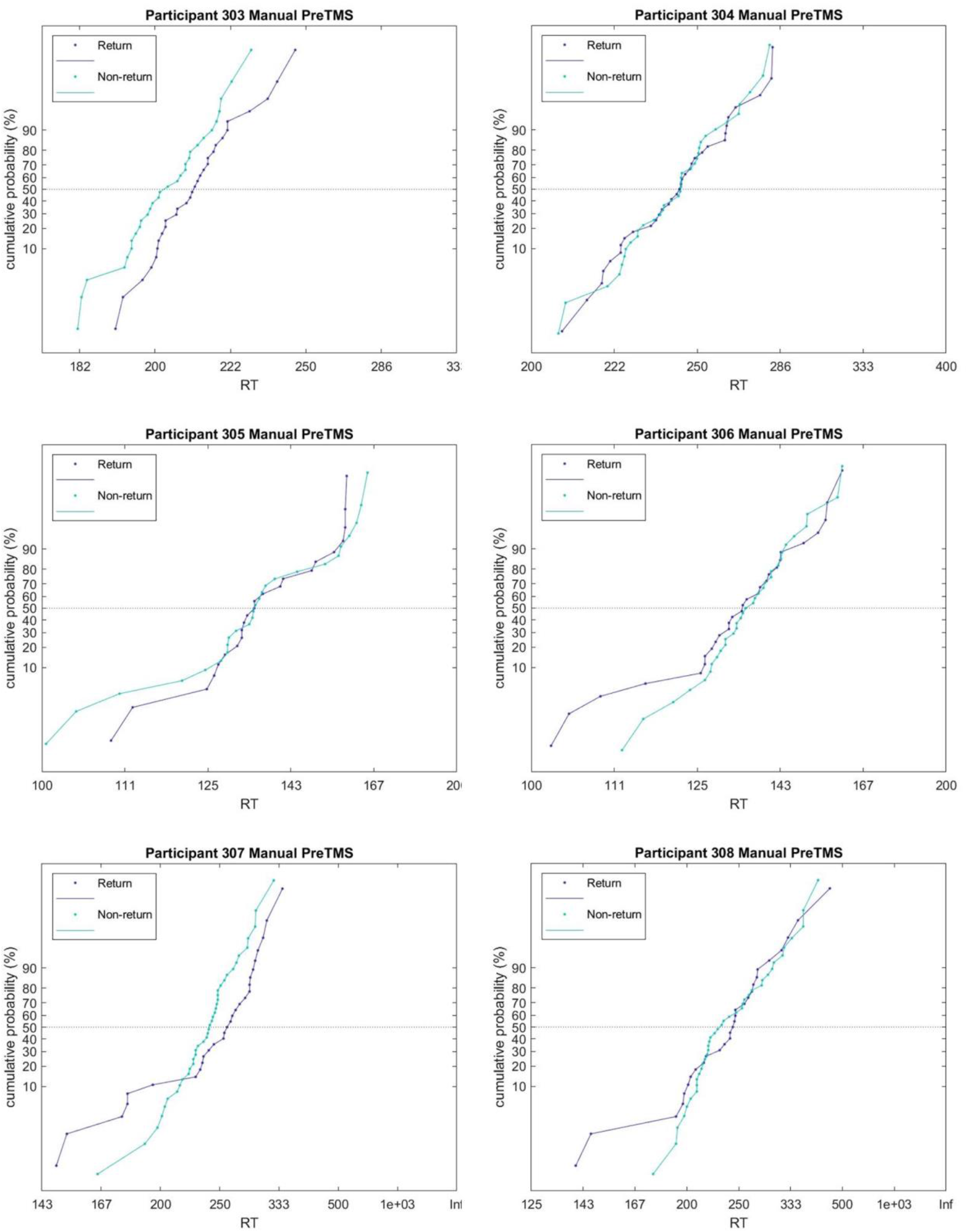
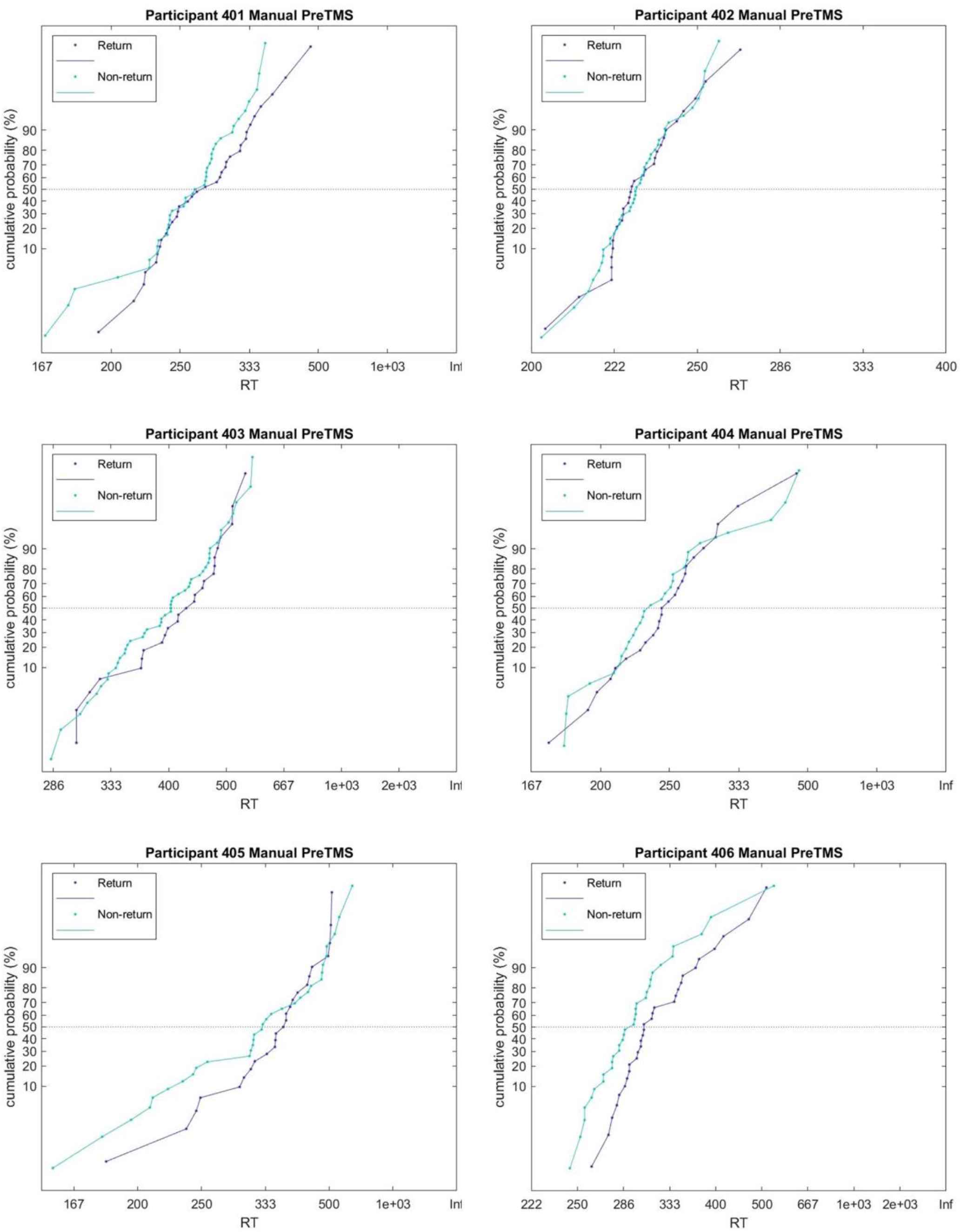
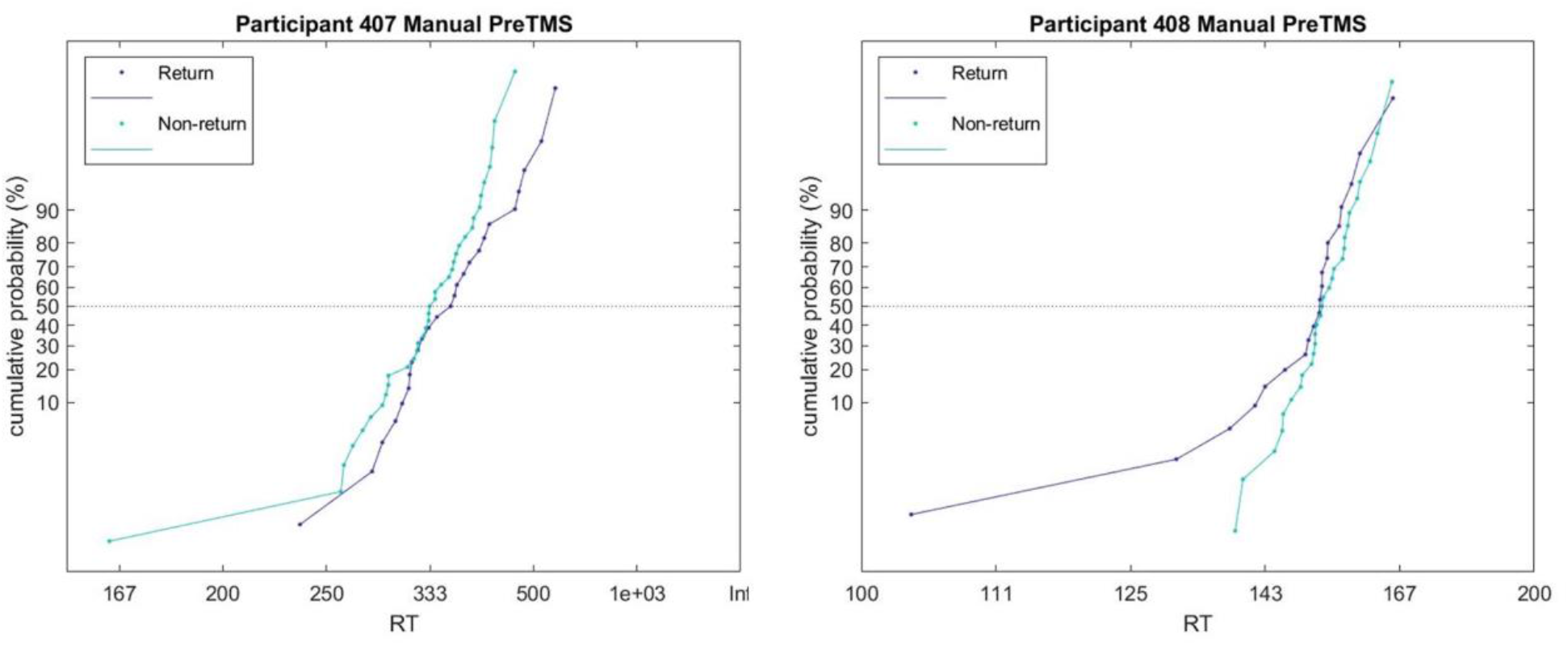
Participants’ individual cumulative RT distributions in the Return condition (grey lines) and Non-return condition (green lines), for Manual IOR.

**Supplementary Figure A2.**
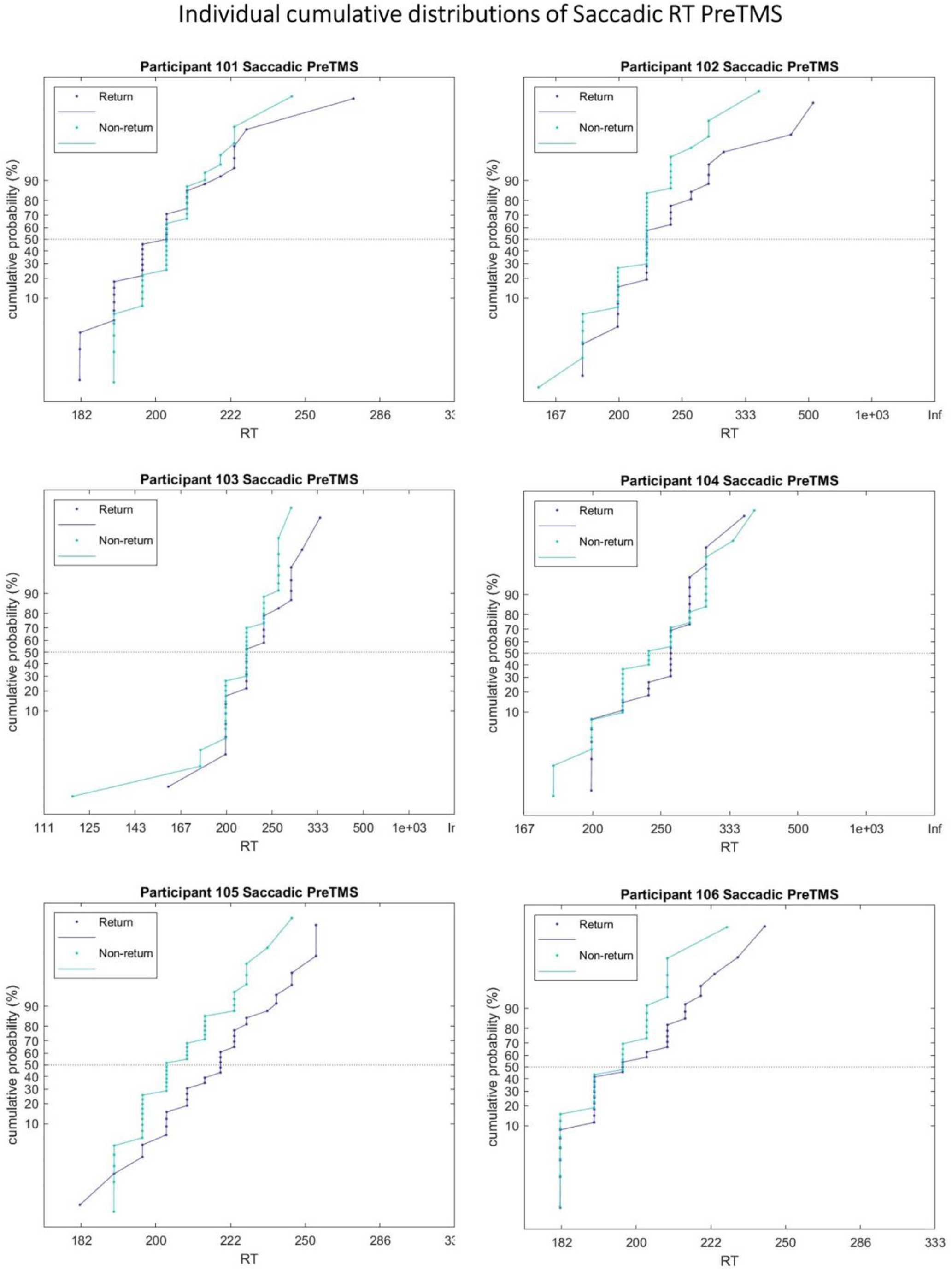
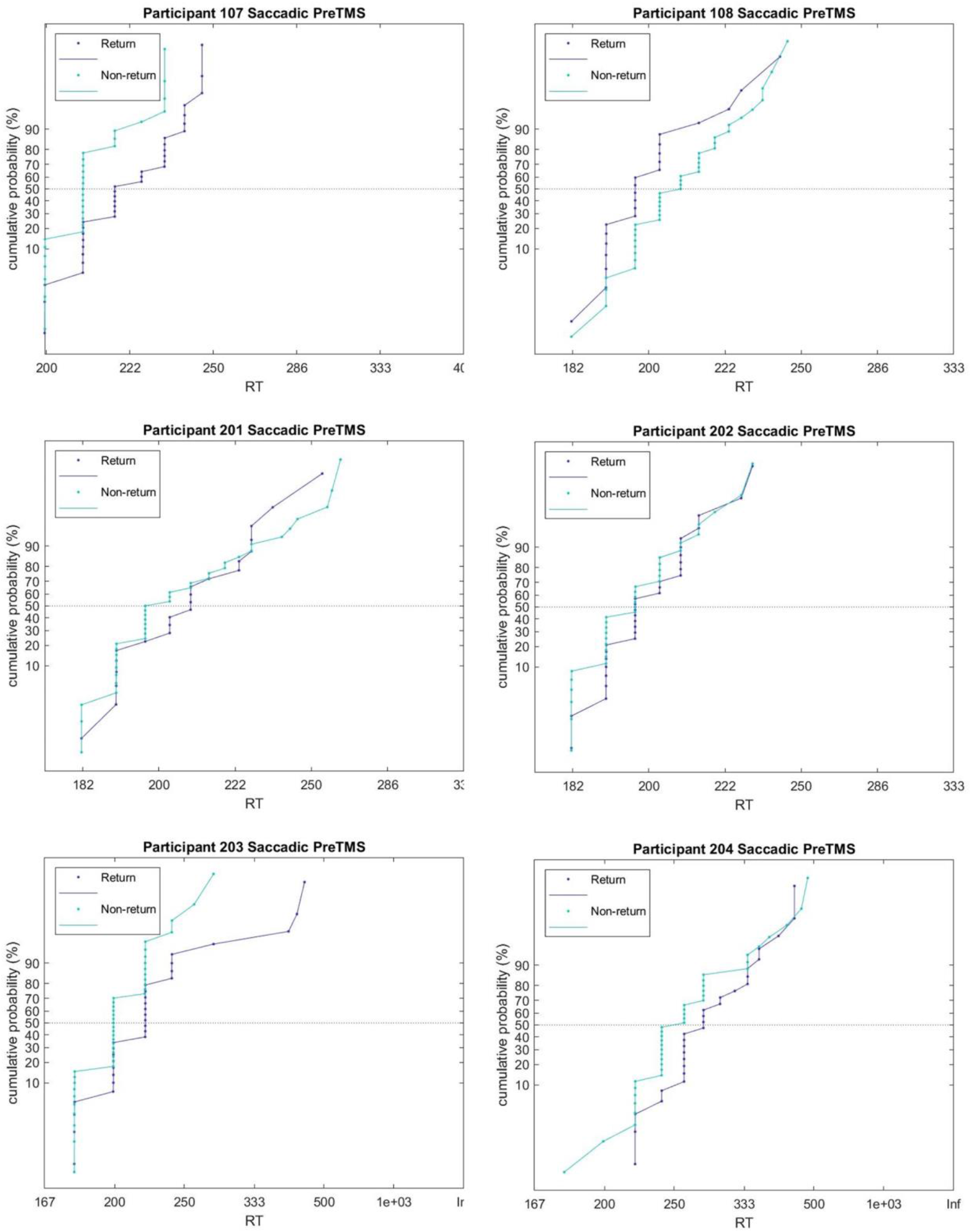
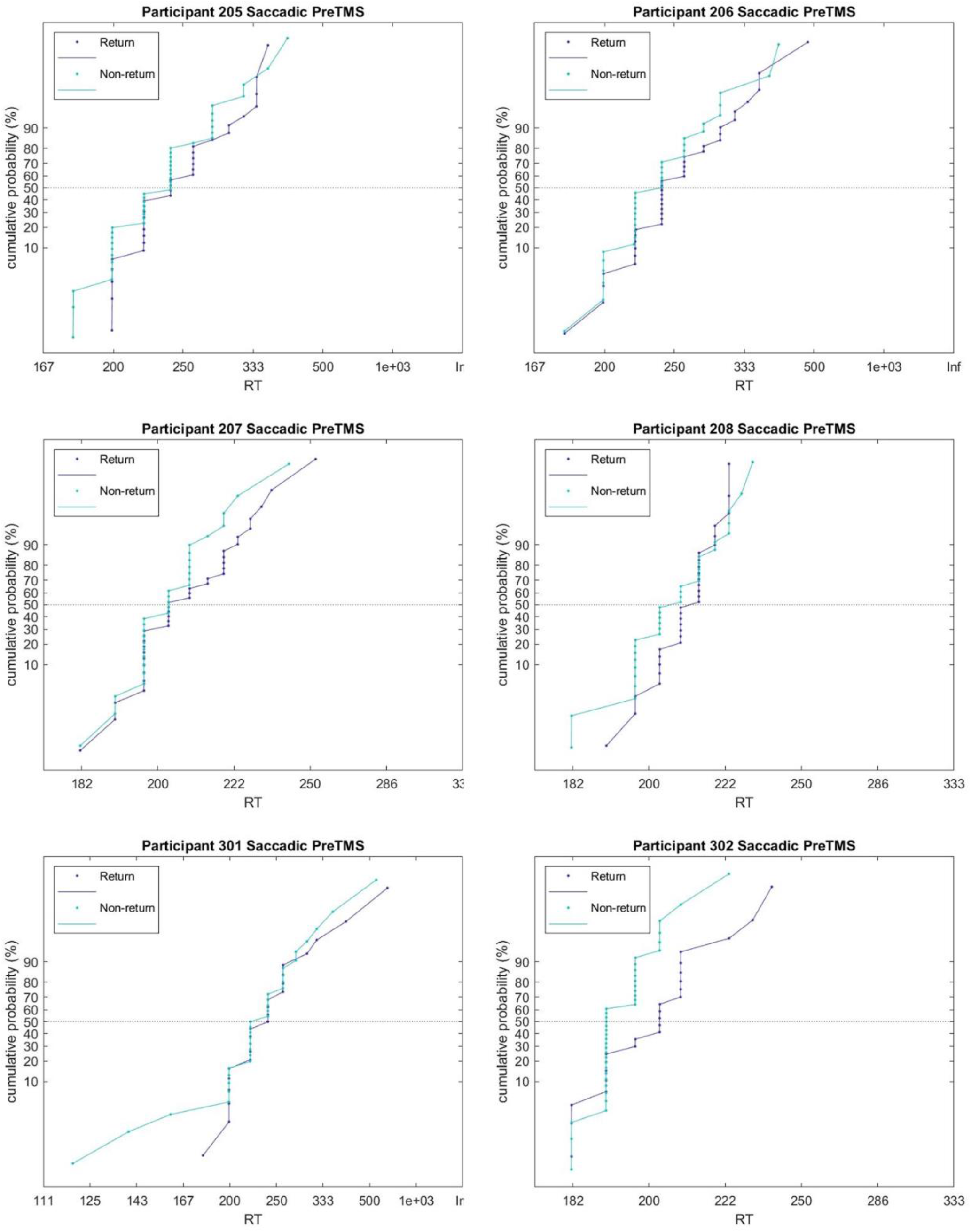
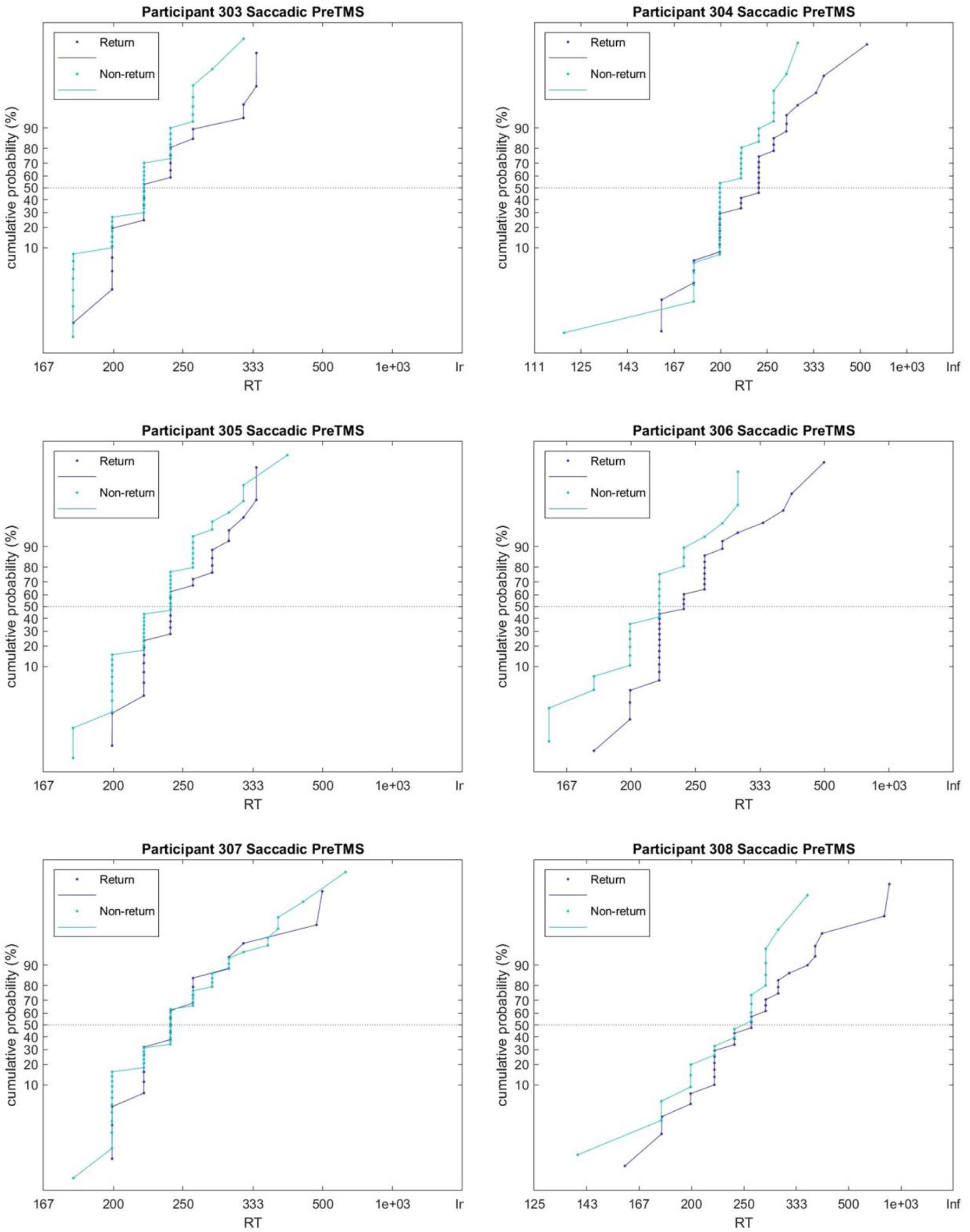
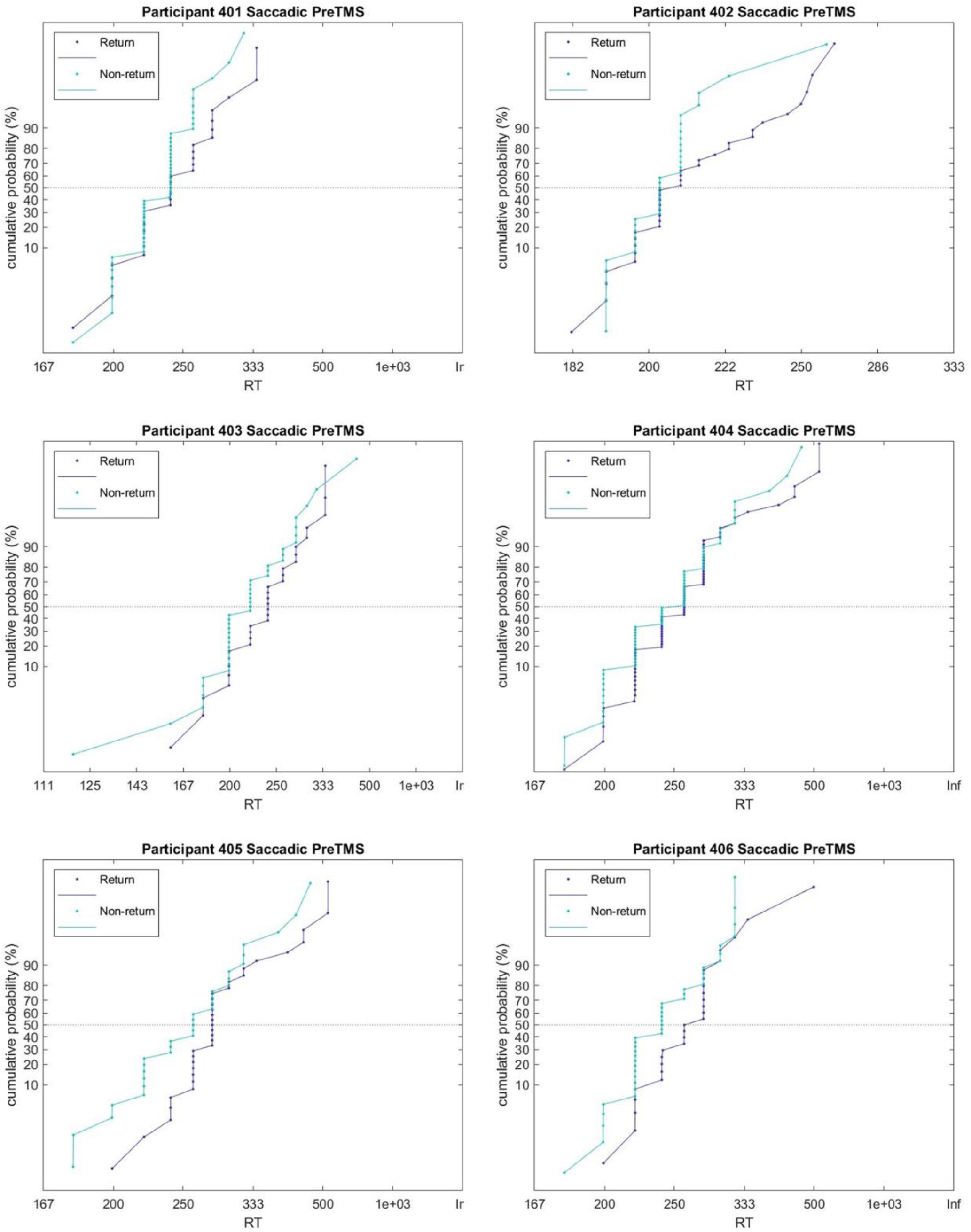
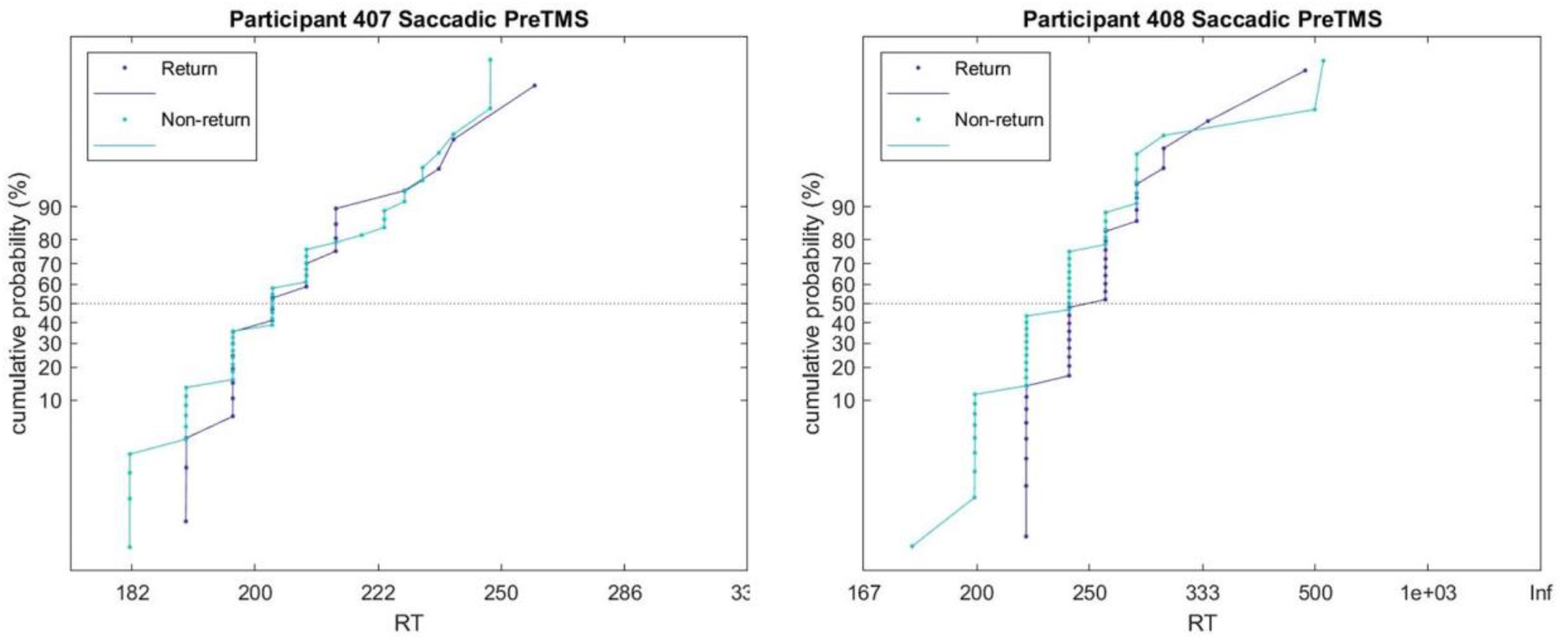
Participants’ individual cumulative RT distributions in the Return condition (grey lines) and Non-return condition (green lines), for Saccadic IOR.

**Supplementary Table A1.**

| Symbol | Parameter | Free* / Fixed |
| --- | --- | --- |
| $\mu$ | Evidence accumulation rate | Free* |
| $\sigma$ | Within-trial noise | Free* |
| $\vartheta$ | Decision threshold | Free* |
| $Da_{go}$ | Non-decision time | Fixed |

\*while the other parameters are fixed (see Methods for details)

## References

Bartolomeo, P., Chokron, S., & Siéroff, E. (1999). Facilitation instead of inhibition for repeated right-sided events in left neglect. Neuroreport, 10(16), 3353–3357.

Bartolomeo, P., & Seidel Malkinson, T. (2019). Hemispheric lateralization of attention processes in the human brain. Current Opinion in Psychology, 29, 90–96.

Bartolomeo, P., Siéroff, E., Decaix, C., & Chokron, S. (2001). Modulating the attentional bias in unilateral neglect: The effects of the strategic set. Experimental Brain Research. Experimentelle Hirnforschung. Experimentation Cerebrale, 137(3/4), 424–431.

Berlucchi, G. (2006). Inhibition of return: a phenomenon in search of a mechanism and a better name. Cognitive Neuropsychology, 23(7), 1065–1074.

Berlucchi, G., Di Stefano, M., Marzi, C. A., Morelli, M., & Tassinari, G. (1981). Direction of attention in the visual field as measured by a reaction time paradigm. Behavioural Brain Research, 2(2), 244–245.

Bisley, J. W., & Goldberg, M. E. (2010). Attention, intention, and priority in the parietal lobe. Annual Review of Neuroscience, 33, 1.

Bompas, A., Hedge, C., & Sumner, P. (2017). Speeded saccadic and manual visuo-motor decisions: Distinct processes but same principles. Cognitive Psychology, 94, 26–52.

Boucher, L., Palmeri, T. J., Logan, G. D., & Schall, J. D. (2007). Inhibitory control in mind and brain: an interactive race model of countermanding saccades. Psychological Review, 114 (2), 376–397.

Bourgeois, A., Chica, A. B., Migliaccio, R., Thiebaut de Schotten, M., & Bartolomeo, P. (2012). Cortical control of inhibition of return: evidence from patients with inferior parietal damage and visual neglect. Neuropsychologia, 50(5), 800–809.

Bourgeois, A., Chica, A. B., Valero-Cabre, A., & Bartolomeo, P. (2013a). Cortical control of inhibition of return: Causal evidence for task-dependent modulations by dorsal and ventral parietal regions. Cortex; a Journal Devoted to the Study of the Nervous System and Behavior, 49(8), 2229–2238.

Bourgeois, A., Chica, A. B., Valero-Cabre, A., & Bartolomeo, P. (2013b). Cortical control of Inhibition of Return: Exploring the causal contributions of the left parietal cortex. Cortex; a Journal Devoted to the Study of the Nervous System and Behavior, 49(10), 2927–2934.

Brown, S. D., & Heathcote, A. (2008). The simplest complete model of choice response time: linear ballistic accumulation. Cognitive Psychology, 57(3), 153–178.

Carpenter, R. H. S., Reddi, B. A. J., & Anderson, A. J. (2009). A simple two-stage model predicts response time distributions. The Journal of Physiology, 587(Pt 16), 4051-4062.

Carpenter, R. H., & Williams, M. L. (1995). Neural computation of log likelihood in control of saccadic eye movements. Nature, 377(6544), 59–62.

Chica, A. B., Bartolomeo, P., & Valero-Cabré, A. (2011). Dorsal and ventral parietal contributions to spatial orienting in the human brain. Journal of Neuroscience, 31(22), 8143–8149.

Chica, A. B., MartÍn-Arévalo, E., Botta, F., & Lupiánez, J. (2014). The Spatial Orienting paradigm: How to design and interpret spatial attention experiments. Neuroscience and Biobehavioral Reviews, 40, 35–51.

Chica, A. B., Taylor, T. L., Lupiáñez, J., & Klein, R. M. (2010). Two mechanisms underlying inhibition of return. Experimental Brain Research, 201(1), 25–35.

Coles, M. G. (1997). Neurons and reaction times. Science, 275(5297), 142–143; author reply 144-145.

Dorris, M. C., Klein, R. M., Everling, S., & Munoz, D. P. (2002). Contribution of the primate superior colliculus to inhibition of return. Journal of Cognitive Neuroscience, 14(8), 1256–1263.

Dorris, M. C., & Munoz, D. P. (1998). Saccadic probability influences motor preparation signals and time to saccadic initiation. The Journal of Neuroscience: The Official Journal of the Society for Neuroscience, 18(17), 7015–7026.

Everling, S., & Munoz, D. P. (2000). Neuronal correlates for preparatory set associated with pro-saccades and anti-saccades in the primate frontal eye field. The Journal of Neuroscience: The Official Journal of the Society for Neuroscience, 20(1), 387–400.

Gold, J. I., & Shadlen, M. N. (2007). The neural basis of decision making. Annual Review of Neuroscience, 30, 535–574.

Hanes, D. P., & Schall, J. D. (1996). Neural control of voluntary movement initiation. Science (New York, N.Y.), 274(5286), 427–430.

Heinen, K., Feredoes, E., Ruff, C. C., & Driver, J. (2017). Functional connectivity between prefrontal and parietal cortex drives visuo-spatial attention shifts. Neuropsychologia, 99, 81–91.

Hilchey, M. D., Klein, R. M., & Satel, J. (2014). Returning to “inhibition of return” by dissociating longterm oculomotor IOR from short-term sensory adaptation and other nonoculomotor “inhibitory” cueing effects. Journal of Experimental Psychology. Human Perception and Performance, 40(4), 1603–1616.

Ivanoff, J., & Klein, R. M. (2001). Attending, intending, and the importance of task settings. In Behavioral and Brain Sciences (Vol. 24, Issue 5, pp. 889–890). 10.1017/s0140525xQ1310105

Jasp Team. (2020). JASP (Version 0.12)[Computer software],

Klein, R. M. (1988). Inhibitory tagging system facilitates visual search. Nature, 334(6181), 430–431.

Klein, R. M. (2000). Inhibition of return. Trends in Cognitive Sciences, 4(4), 138–147.

Klein, R. M., Dölek, S., & Christie, J. (n.d.). The output form of inhibition of return likely operates at and after the bottleneck. Acta Psychologica.

Klein, R. M., Kavyani, M., Farsi, A., & Lawrence, M. A. (2020). Using the locus of slack logic to determine whether the output form of inhibition of return affects an early or late stage of processing. Cortex; a Journal Devoted to the Study of the Nervous System and Behavior, 122, 123–130.

Klein, R. M., & Redden, R. S. (2018). Two “inhibitions of return” bias orienting differently. Spatial Biases in Perception and Cognition, 295–306.

Klein, R. M., Redden, R. S., & Hilchey, M. D. (2023). Visual search and the inhibitions of return. Frontiers in Cognition, 2. 10.3389/fcogn.2023.1146511

Lev-Ari, T., Zahar, Y., Agarwal, A., & Gutfreund, Y. (2020). Behavioral and neuronal study of inhibition of return in barn owls. Scientific Reports, 10(1), 7267.

Lisi, M., & Cavanagh, P. (2015). Dissociation between the Perceptual and Saccadic Localization of Moving Objects. Current Biology: CB, 25(19), 2535–2540.

Lisi, M., & Cavanagh, P. (2017). Different spatial representations guide eye and hand movements. Journal of Vision, 17(2), 12.

Luce, D. R. (1986). Response Times: Their Role in Inferring Elementary Mental Organization. OUP USA.

Ludwig, C. J. H., Farrell, S., Ellis, L. A., & Gilchrist, I. D. (2009). The mechanism underlying inhibition of saccadic return. Cognitive Psychology, 59(2), 180–202.

Lupiáñez, J. (2010). Inhibition of return. Attention and Time, 17–34.

Lupiáñez, J., Decaix, C., Siéroff, E., Chokron, S., Milliken, B., & Bartolomeo, P. (2004). Independent effects of endogenous and exogenous spatial cueing: inhibition of return at endogenously attended target locations. Experimental Brain Research. Experimentelle Hirnforschung. Experimentation Cerebrale, 159(4), 447–457.

Lupiáñez, J., Klein, R. M., & Bartolomeo, P. (2006). Inhibition of return: Twenty years after. Cognitive Neuropsychology, 23(7), 1003–1014.

Maclnnes, W. J. (2017). Multiple Diffusion Models to Compare Saccadic and Manual Responses for Inhibition of Return. Neural Computation, 29(3), 804–824.

MATLAB. (2017). version R2017b (Version 2017b). The MathWorks Inc.

Maylor, E. A., & Hockey, R. (1985). Inhibitory component of externally controlled covert orienting in visual space. Journal of Experimental Psychology. Human Perception and Performance, 11(6), 777787.

McDonald, J. J., Ward, L. M., & Kiehl, K. A. (1999). An event-related brain potential study of inhibition of return. Perception & Psychophysics, 61(7), 1411–1423.

Miller, I. K., Gochin, P. M., & Gross, C. G. (1991). Habituation-like decrease in the responses of neurons in inferior temporal cortex of the macaque. In Visual Neuroscience (Vol. 7, Issue 4, pp. 357–362). 10.1017/s09525238Q0004843

Mirpour, K., Arcizet, F., Ong, W. S., & Bisley, J. W. (2009). Been there, seen that: a neural mechanism for performing efficient visual search. Journal of Neurophysiology, 102(6), 3481–3491.

Mirpour, K., Bolandnazar, Z., & Bisley, J. W. (2019). Neurons in FEF Keep Track of Items That Have Been Previously Fixated in Free Viewing Visual Search. The Journal of Neuroscience: The Official Journal of the Society for Neuroscience, 39(11), 2114–2124.

Munoz, D. P., & Everling, S. (2004). Look away: the anti-saccade task and the voluntary control of eye movement. Nature Reviews. Neuroscience, 5(3), 218–228.

Mushquash, A. R., Fawcett, J. M., & Klein, R. M. (2012). Inhibition of return and schizophrenia: a metaanalysis. Schizophrenia Research, 135(1-3), 55–61.

Patel, S. S., Peng, X., & Sereno, A. B. (2010). Shape effects on reflexive spatial selective attention and a plausible neurophysiological model. Vision Research, 50(13), 1235–1248.

Posner, M. I., & Cohen, Y. (1984). Components of visual orienting. In H. Bouma & D. Bouwhuis (Eds.), Attention and performance X: Control of language processes (pp. 531–556.). Erlbaum.

Posner, M. I., Rafal, R. D., Choate, L. S., & Vaughan, J. (1985). Inhibition of return: Neural basis and function. Cognitive Neuropsychology, 2, 211–228.

Pouget, P., Emeric, E. E., Stuphorn, V., Reis, K., & Schall, J. D. (2005). Chronometry of visual responses in frontal eye field, supplementary eye field, and anterior cingulate cortex. Journal of Neurophysiology, 94(3), 2086–2092.

Pouget, P., Logan, G. D., Palmeri, T. J., Boucher, L., Paré, M., & Schall, J. D. (2011). Neural basis of adaptive response time adjustment during saccade countermanding. The Journal of Neuroscience: The Official Journal of the Society for Neuroscience, 31(35), 12604–12612.

Pouget, P., Stepniewska, I., Crowder, E. A., Leslie, M. W., Emeric, E. E., Nelson, M. J., & Schall, J. D. (2009). Visual and motor connectivity and the distribution of calcium-binding proteins in macaque frontal eye field: implications for saccade target selection. Frontiers in Neuroanatomy, 3, 2.

Purcell, B. A., Schall, J. D., Logan, G. D., & Palmeri, T. J. (2012). From Salience to Saccades: MultipleAlternative Gated Stochastic Accumulator Model of Visual Search. In Journal of Neuroscience (Vol. 32, Issue 10, pp. 3433–3446). 10.1523/jneurosci.4622-ll.2012

Rafal, R., Egly, R., & Rhodes, D. (1994). Effects of inhibition of return on voluntary and visually guided saccades. Canadian Journal of Experimental Psychology = Revue Canadienne de Psychologie Experimentale, 48(2), 284–300.

Ratcliff, R., Hasegawa, Y. T., Hasegawa, R. P., Smith, P. L., & Segraves, M. A. (2007). Dual diffusion model for single-cell recording data from the superior colliculus in a brightness-discrimination task. Journal of Neurophysiology, 97(2), 1756–1774.

Redden, R. S., Hilchey, M. D., & Klein, R. M. (2016). Peripheral stimuli generate different forms of inhibition of return when participants make prosaccades versus antisaccades to them. Attention, Perception & Psychophysics, 78(8), 2283–2291.

Redden, R. S., MacInnes, W. J., & Klein, R. M. (2020). Inhibition of return: An information processing theory of its natures and significance. Cortex; a Journal Devoted to the Study of the Nervous System and Behavior. 10.1016/i.cortex.2020.ll.009

Sapir, A., Soroker, N., Berger, A., & Henik, A. (1999). Inhibition of return in spatial attention: direct evidence for collicular generation. Nature Neuroscience, 2(12), 1053–1054.

Satel, J., Wilson, N. R., & Klein, R. M. (2019). What Neuroscientific Studies Tell Us about Inhibition of Return. Vision (Basel, Switzerland), 3(4). 10.3390/vision3040Q58

Schall, J. D. (2004). On the role of frontal eye field in guiding attention and saccades. Vision Research, 44(12), 1453–1467.

Schall, J. D. (2019). Accumulators, Neurons, and Response Time. Trends in Neurosciences, 42(12), 848–860.

Schall, J. D., & Hanes, D. P. (1993). Neural basis of saccade target selection in frontal eye field during visual search. Nature, 366(6454), 467–469.

Seidel Malkinson, T., & Bartolomeo, P. (2018). Fronto-parietal organization for response times in inhibition of return: The FORTIOR model. Cortex; a Journal Devoted to the Study of the Nervous System and Behavior, 102, 176–192.

Seidel Malkinson, T., Bayle, D. J., Kaufmann, B. C., Liu, J., Bourgeois, A., Lehongre, K., Fernandez-Vidal, S., Navarro, V., Lambrecq, V., Adam, C., Margulies, D. S., Sitt, J. D., & Bartolomeo, P. (2024). Intracortical recordings reveal vision-to-action cortical gradients driving human exogenous attention. Nature Communications, 15(1), 2586.

Smith, P. L., & Ratcliff, R. (2004). Psychology and neurobiology of simple decisions. Trends in Neurosciences, 27(3), 161–168.

Spagna, A., Liu, J., & Bartolomeo, P. (2026). Beta and gamma dynamics in attentional networks predict conscious reports. The Journal of Neuroscience: The Official Journal of the Society for Neuroscience, 60590252026.

Steinmetz, M. A., Connor, C. E., Constantinidis, C., & McLaughlin, J. R. (1994). Covert attention suppresses neuronal responses in area 7a of the posterior parietal cortex. Journal of Neurophysiology, 72(2), 1020–1023.

Steinmetz, M. A., & Constantinidis, C. (1995). Neurophysiological evidence for a role of posterior parietal cortex in redirecting visual attention. Cerebral Cortex, 5(5), 448–456.

Taylor, T. L., & Klein, R. M. (2000). Visual and motor effects in inhibition of return. Journal of Experimental Psychology. Human Perception and Performance, 26(5), 1639–1656.

Vivas, A. B., Humphreys, G. W., & Fuentes, L. J. (2006). Abnormal inhibition of return: A review and new data on patients with parietal lobe damage. Cognitive Neuropsychology, 23(7), 1049–1064.

